# Interpretable Metric Learning in Comparative Metagenomics: The Adaptive Haar-like Distance

**DOI:** 10.1101/2023.09.27.559681

**Authors:** Evan Gorman, Manuel E. Lladser

## Abstract

Random forests have emerged as a promising tool in comparative metagenomics because they can predict environmental characteristics based on microbial composition in datasets where *β*-diversity metrics fall short of revealing meaningful relationships between samples. Nevertheless, despite this efficacy, they lack biological insight in tandem with their predictions, potentially hindering scientific advancement. To overcome this limitation, we leverage a geometric characterization of random forests to introduce a data-driven phylogenetic *β*-diversity metric, the adaptive Haar-like distance. This new metric assigns a weight to each internal node (i.e., split or bifurcation) of a reference phylogeny, indicating the relative importance of that node in discerning environmental samples based on their microbial composition. Alongside this, a weighted nearest-neighbors classifier, constructed using the adaptive metric, can be used as a proxy for the random forest while maintaining accuracy on par with that of the original forest and another state-of-the-art classifier, CoDaCoRe. As shown in datasets from diverse microbial environments, however, the new metric and classifier significantly enhance the biological interpretability and visualization of high-dimensional metagenomic samples.

**Author summary:** Traditional phylogenetic *β*-diversity metrics, particularly weighted and unweighted UniFrac, have had great success in comparing and visualizing high-dimensional metagenomic samples. Nonetheless, these metrics rely upon pre-established biological assumptions that might not capture key microbial players or relationships between some samples. On the contrary, supervised machine learning algorithms, such as random forests, can often capture intricate relationships between microbial samples; however, unveiling these relationships is often challenging due to the intricate inner mechanisms inherent to these algorithms.

The adaptive Haar-like distance integrates the merits of *β*-diversity metrics and random forests, allowing for precise, intuitive, and visual comparison of metagenomic samples, offering valuable scientific insight into the distinctions and associations among microbial environments.

## Introduction

Comparative metagenomics seeks to identify conserved or variable genetic features across microbial communities to discern the relationship between environmental characteristics and microbial composition. Phylogenetic *β*-diversity metrics facilitate this process using a reference phylogenetic tree. The standard analysis involves a pipeline that clusters sequence reads (obtained from one or multiple environments) into Operational Taxonomic Units (OTUs) based on a predetermined level of sequence similarity. The OTUs are then consolidated into feature tables with abundance counts per sample. A common practice is to map these OTUs onto the leaves of a reference phylogenetic tree such as Greengenes [1], Silva [2], or WoL [3]. If a high resolution of sequence similarity was used to define the OTUs, such as the conventional 97% or 99%, the processed data can have hundreds of thousands of dimensions, which poses significant challenges for its analysis.

Phylogenetic *β*-diversity metrics assume that OTUs with shared evolutionary histories possess similar traits, which may be advantageous or disadvantageous in environments with comparable characteristics; in particular, samples containing closely related OTUs should exhibit closer clustering. These metrics are commonly employed to assess the significance of clustering or correlation with covariates such as pH, salinity, or depth [4], among many others. Phylogenetic *β*-diversity metrics are also often used alongside principle coordinates analysis (PCoA) [5] to generate low-dimensional visualizations of microbial datasets [6].

UniFrac [7] is arguably the most renowned phylogenetic *β*-diversity metric. Its fundamental breakthrough lies in effectively integrating microbes’ phylogenetic relatedness: differences in OTU composition between environments are weighted by the shared length of evolutionary history among the OTUs. This metric has two variants, weighted and unweighted. Weighted UniFrac can be viewed as an Earth Mover’s distance, where the ground metric is defined by the underlying reference phylogeny [8].

Double Principle Coordinates Analysis (DPCoA) [9] is another, albeit less well-known, *β*-diversity metric. It is a Mahalanobis-type distance [10] associated with the inverse of the so-called phylogenetic covariance matrix of the reference phylogenetic tree (see definitions 1.2 and 1.3). This matrix encodes the shared branch length, leading to the root, between all pairs of OTUs [11]. In particular, the entries of the phylogenetic covariance matrix can be interpreted as the covariance of a specific trait that evolves over the reference phylogenetic according to a Brownian motion [12, Chapter 3].

DPCoA can be considered a Euclidean version (aka *ℓ*^2^-version) of weighted UniFrac, as both metrics rely on the same fundamental assumptions regarding OTU relatedness. (Conversely, UniFrac can be seen as an *ℓ*^1^-version of DPCoA.) While UniFrac and DPCoA and their associated embeddings have demonstrated remarkable efficacy across diverse microbial scenarios, explaining their embeddings solely based on microbial abundances remains a challenge.

Recent work showed that discrete Haar-like wavelets could significantly pseudodiagonalize (i.e., sparsify) the phylogenetic covariance matrix (via a change of basis) of most large binary trees, which motivated the introduction of a novel phylogenetic *β*-diversity metric known as the Haar-like distance [13]. This new metric may be regarded as a proxy for DPCoA; however, unlike DPCoA and UniFrac, it admits a simple decomposition in terms of the splits or bifurcations (i.e., internal nodes) of the reference phylogeny, which enables further interpretation and visualization of microbial sample distances in terms of differences between microbial clade abundances. Despite its breakthrough, the Haar-like distance, along with all existing phylogenetic *β*-diversity metrics, is also constrained by the inherent assumptions in the reference phylogeny (specifically, encoded in its covariance matrix). This bears the question, *can these metrics be tailored to capture more subtle relationships within specific metagenomic datasets, similar to those offered by supervised machine learning techniques, while still leveraging the evolutionary relationships that β-diversity metrics have successfully exploited?*

Random forests (RFs) are a popular supervised machine learning method that combine multiple decision trees to make predictions [14]. While specific architectures may vary among implementations, the fundamental idea is as follows: each tree is built on a random subset of labeled training data and optimizes a criterion such as Gini impurity [15] to cluster data with similar labels. To make predictions on new unlabeled data, each tree returns a label, and the RF prediction is based on the (potentially weighted) average of these individual tree predictions.

RFs usually exhibit superior or comparative performance to other state-of-the-art methods in microbiome host-trait prediction [16], and numerous studies have documented their effectiveness in metagenomics classification tasks [17–22]. While these classifiers achieve impressive accuracy through the averaging of (random) decision trees, the inherent randomness in their construction can obscure the learned relationships between OTU composition and prediction. Furthermore, RF predictions cannot be explained solely by existing feature importance measures, such as Gini or permutation importance, as they can be highly sensitive to correlated or highly variable features [23, 24].

In this manuscript, we introduce a new phylogenetic *β*-diversity metric: the adaptive Haar-like distance, which is inspired by the recent Haar-like distance [13]. Taking a metric learning approach [25], our algorithm learns a data-dependent weighting of the most important phylogenetic relationships across a set of samples to discover robust representations of microbial abundance patterns. In contrast to traditional metric learning algorithms, which are known to be computationally expensive [25] and suffer from the curse of dimensionality, our approach scales well with large datasets.

Scalability is achieved by leveraging a pretrained RF classifier, which we adapt to fit a metric: a Haar-like distance associated with a phylogenetic covariance matrix that we learn from the classifier. Accordingly, the adaptive Haar-like distance combines the predictive power of RFs with the interpretability of the Haar-like distance.

## Materials and Methods

In this section, we outline our metric learning algorithm. First, we discuss the Haar-like wavelet basis and its corresponding coordinate system, which gives rise to the Haar-like distance [13]. We then generalize this metric by introducing tunable weight parameters, leading to the adaptive Haar-like distance and corresponding kernel.

Next, to learn weights in a data-dependent manner, we examine the representation of random forests as local average estimators [26]. Here, random forest classifiers are framed as kernel estimators built from their so-called affinity.

Finally, we use a compressed sensing algorithm to learn a sparse set of weights to approximate a random forest affinity by an adaptive Haar-like kernel. This substitution yields a surrogate random forest model that is precisely interpretable through a limited number of Haar-like coordinates, representing the most relevant clade abundances for distinguishing between environments in a given dataset.

### Haar-like Wavelet Basis

The Haar-like wavelets were first described in [27] for the multiscale analysis of datasets equipped with a hierarchical partition tree. The wavelets form an orthonormal basis for the vector space of (real-valued) functions defined on the leaves of such trees and localize information on the leaves at scales determined by the proximity of each internal node to the (external) root: the closer an internal node is to the root, the coarser the scale associated with that node.

In phylogenetic trees (particularly out-rooted, see [13, Definition 2.1]), there is a direct correspondence between the Haar-like wavelets and the internal nodes. In particular, since the latter represent speciation events that group OTUs into clades, the Haar-like wavelets offer a basis for comparing clades of microorganisms as opposed to separate OTUs. So, assuming sample abundances correlate within the same clade across similar environments, projecting these onto the wavelets should elucidate relationships between microbial composition and environmental factors [13].

### Haar-like Coordinates

Before describing how to project functions defined over the leaves of a phylogenetic tree onto its Haar-like wavelet basis, we introduce some notation. In what follows, *T* denotes a reference phylogenetic tree with vertex and edge set *V* and *E*, respectively, and branch length function *ℓ* : *E*→ [0, + ∞). In practice, the leaves of *T* represent OTUs, whereas its interior nodes represent inferred speciation events.

We distinguish the set of leaves *L* from the set of internal nodes *I*, noting that they partition *V* (i.e., *L* ∪ *I* = *V* but *L* ∩ *I* = ∅). The root of *T* is labeled by ○. For any internal node *v* ∈ *I*, denote by *L*(*v*) the set of leaves that descend from *v*. Further, if *v* has at least two children, *v*_+_ and *v*_*−*_ denote the left and right children descending from it, respectively. (In [13], these were denoted *v*0 and *v*1, respectively.)

We assume that microbial abundance data on the leaves of *T* are normalized to sum to one so that each sample can be represented as a probability mass function on *L*. We denote these functions by *x, x*_1_, *x*_2_, …; in particular, *x* : *L* → [0, 1) satisfies that ∑_*v∈L*_ *x*(*v*) = 1. Therefore, each sample is compositional (that is, distribution valued) and could be analyzed using a variety of methods [28].

With the above notation, the projection of a sample *x* onto a Haar-like wavelet *φ*_*v*_, associated with internal node *v* ≠ ○, can be conveniently represented in terms of average abundances on subtrees of the reference phylogeny.

**Definition 1.1** (Average). For a given function *x* : *L* → ℝ and non-empty *J* ⊂ *L* of cardinality |*J* |, we define the mean of *x* over *J* as

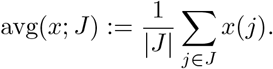

**Theorem 1.1** (Projection onto *φ*_*v*_). *Let v≠* ○ *be an interior node of T*. *The projection of a function x* : *L* → ℝ *onto Haar-like wavelet φ*_*v*_ *is:*

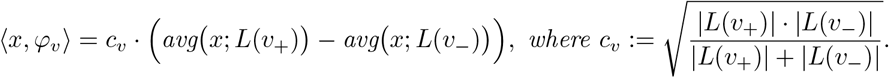

We refer to the set of projections {⟨*x, φ*_*v*_⟩}_*v∈I\{○}*_ as the **Haar-like coordinates** associated with a sample *x*. (We disregard the root of *T* in our setting because, for compositional data *x*,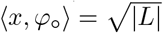, i.e. a constant. In particular, as we assess microbial samples by differences in their Haar-like coordinates, this coordinate holds no relevance in our framework.)

We highlight that the Haar-like coordinates of log(*x*) (i.e., the function log *x*(*v*) when *x* ∈ (*v*) *>* 0 for all *v* ∈ *L*) correspond to the isometric log-ratio (ILR) coordinates [29], which have been used in previous metagenomics analyses like PhILR [30] and Phylofactorization [31]. The ILR coordinates necessitate zero-count replacement and employ logarithmic ratios of geometric means to map compositional data into an unconstrained Hilbert space, known as the Bayes space [32], where the Euclidean distance is replaced by the Aitchison distance [33]. This distance is invariant to the underlying phylogenetic structure and thus loses the assumptions of evolutionary relatedness that have been key to the success of phylogenetic *β*-diversity metrics.

### Haar-like Distance

As mentioned earlier, DPCoA is a Mahalanobis-type distance associated with the inverse of the phylogenetic covariance matrix of the reference tree. The precise interpretations of this statement follow.

**Definition 1.2** (Phylogenetic Covariance). For nodes *i, j* ∈ *V*, let [*i, j*] denote the set of edges in the shortest path between nodes *i* and *j* in *T*. Also, let (*i* ∧ *j*) be the least common ancestor of *i* and *j*. Namely, the *v* ∈ *V* that maximizes | [*v*, ○] | among all the nodes that are ancestors to both *i* and *j*. The phylogenetic covariance matrix of *T* is the matrix of dimensions |*L*| × |*L*| with entries

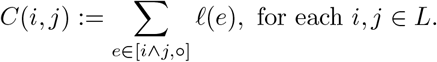

**Definition 1.3** (Double Principal Coordinate Analysis [9]). The DPCoA distance between two environmental samples *x*_1_ and *x*_2_ is

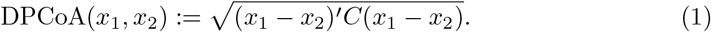

Let Φ denote the matrix whose columns consist of the Haar-like wavelets of the reference phylogeny. On large trees, if one changes basis using Φ, then, in the new coordinates, and with high probability, *C* will be nearly diagonal [13, Corollary 3.8]. Namely, the matrix Φ^*′*^ *C* Φ is significantly sparse, which motivates substituting *C* by the diagonal matrix

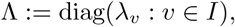

where

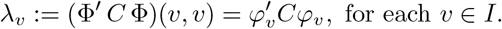

In terms of DPCoA, this is equivalent to substituting the matrix *C* in (1) by the matrix Φ^*′*^ Λ Φ, which motivates the next definition.

**Definition 1.4** (Haar-like Distance [13]). The Haar-like distance between two environmental samples *x*_1_ and *x*_2_ is

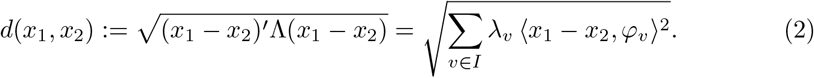

The Haar-like distance is therefore a weighted Euclidean distance between the Haar-like coordinates of pairs of samples. This metric provides an interpretable version of DPCoA, as the distance between two samples relates to a sum indexed by the internal nodes of the reference phylogeny and, unlike the Aitchison distance, includes assumptions about phylogenetic relatedness when calculating distances.

### Adaptive Haar-like Distance and Kernel

Although traditional phylogenetic *β*-diversity metrics have provided significant insights across various datasets, they may not consistently differentiate between samples from distinct environments. On the other hand, despite its biological interpretability, a fundamental limitation of the newly introduced Haar-like distance is the potentially broad biological assumptions encoded by the coefficients *λ*_*v*_, with *v* ∈ *I*, used to define it (see equation (2)). In fact, it seems improbable that these fixed “universal” weights adequately account for relevant differences in microbial composition across arbitrary pairs of environments. Nevertheless, the Haar-like distance allows for easy adjustment of these assumptions by replacing its fixed weights with adaptive ones learned from labeled datasets. We next define this generalization.

**Definition 1.5** (Adaptive Haar-like Kernel & Distance). The Adaptive Haar-like kernel associated with a weight vector *w* = {*w*_*v*_}_*v∈I*_, with *w*_*v*_ *>* 0 for each *v*, is defined as

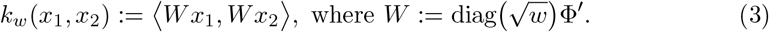

The associated adaptive Haar-like distance between two environmental samples *x*_1_ and *x*_2_ is

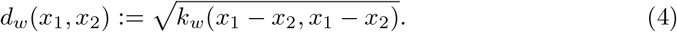

The adaptive Haar-like distance is induced by an inner product between differences of Haar-like coordinates; in particular, it is faithful to the topology of the reference tree. Importantly, each weight aligns with an internal node in the phylogeny, allowing selective weight adjustment for specific clades. This is especially promising for a multiscale analysis of *β*-diversity in sample comparisons. In practice, however, identifying the most important internal nodes in a given context is not immediately clear. The subsequent aim is to choose weights that minimize the adaptive Haar-like distance or maximize the related kernel between samples that share similar environmental characteristics. For efficient weight learning, we turn to random forests (RFs).

### Towards an Interpretable Random Forest Surrogate

Though their connection to metric learning is not immediately clear, an RF can be re-framed as a kernel method by considering the geometry learned through training: each decision tree is a collection of binary decision rules that partition a feature space. By examining the splits made by the decision trees, it is possible to define a notion of similarity between data points based on the trees’ paths they traverse within the forest.

The RF affinity [26] is a kernel that quantifies how often two data points land in the same partition across a forest’s decision trees. This kernel can be used to replicate RF predictions: the label for a new point is estimated through a weighted average of the closest training point labels based on some similarity measure.

In what follows, ⟦·⟧ denotes the indicator function (aka Iverson bracket) of the proposition within, i.e. ⟦· ⟧ = 1 when the statement within parentheses is true; otherwise ⟦· ⟧ = 0.

**Definition 1.6** (Random Forest Affinity [26] & Dissimilarity). Consider an RF consisting of *M* decision trees trained on *n* labeled samples (*x*_*i*_, *y*_*i*_) ∈ (ℝ^*d*^, *𝒞*), where *𝒞* ⊂ ℝ is non-empty. (For example, *𝒞* = {−1, 1} in binary classification, but *𝒞* = ℝ in most regression problems.) For each *x* ∈ ℝ^*d*^, let *ℒ*_*m*_(*x*) denote the leaf (i.e., bin) containing *x* in the *m*-th decision tree. The RF affinity and dissimilarity between two points *x*_1_, *x*_2_ ∈ ℝ^*d*^ are defined, respectively, as follows:

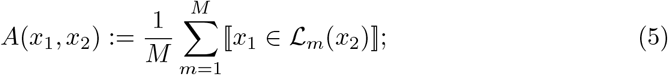

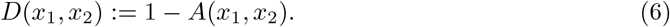

The RF affinity between two points is the fraction of decision trees that group them in the same leaf across the forest; in particular, it is symmetric and measures how similar two points are from the perspective of the trained RF. Accordingly, the dissimilarity is also symmetric but measures how often two training points are placed into different bins by the RF.

In the context of regression, the affinity can be used to construct the so-called kernel RF estimate from labeled samples (*x*_1_, *y*_1_), …, (*x*_*n*_, *y*_*n*_), as follows.

**Definition 1.7** (RFs as Regressive Local Average Estimators [26]). The kernel RF estimate (KeRFE) of a function *f* : ℝ^*d*^ → ℝ at a point *x* is

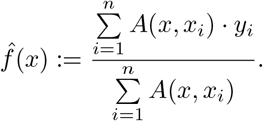

Under mild conditions, the KeRFE converges to the original RF estimate as *n* increases.

KeRFEs do not entirely resolve the interpretability issue of RFs. One reason is the nonstationarity of *A*(*x*_1_, − *x*_2_) [34], as it depends on both *x*_1_ and *x*_2_ rather than just (*x*_1_ *x*_2_), which complicates generating a consistent explanation of the affinity’s behavior across a dataset. This is unlike the adaptive Haar-like kernel and distance, where weights signal the importance of clade abundances in sample comparisons. Nevertheless, as we shall see next, KeRFEs offer a perspective for directly studying the behavior of RFs through their affinity.

### Metric Learning Algorithm

**The core idea in this manuscript** is to learn a weight vector *w* such that the Haar-like kernel (equation (3)) can induce the RF dissimilarity (equation (6)) across the whole training set. In particular, because each weight *w*_*v*_ is directly linked to the internal node *v* and, therefore, a speciation event in the tree, the associated Haar-like kernel may serve as an interpretable proxy for the RF model—from the perspective of the phylogeny. We note that after learning the appropriate weights, the adaptive Haar-like distance and its associated embedding can be recovered from the kernel.

Assume as given *n* labeled samples (*x*_*i*_, *y*_*i*_) ∈ (ℝ^*d*^, ℝ) and collect the *x*_*i*_’s in a data matrix *X* ∈ ℝ^*d×n*^. To accomplish our goal, we first train an RF on the data and recover a pairwise affinity matrix *A* ∈ ℝ^*n×n*^ and dissimilarity matrix *D* := **11**^*′*^ − *A*, where **1** is a column vector of ones of dimension *n*. (The entry in row-*x*_*i*_ and column-*x*_*j*_ of *A* and *D* are *A*(*x*_*i*_, *x*_*j*_) and *D*(*x*_*i*_, *x*_*j*_), respectively.)

Although *D* is in general non-Euclidean, we can use principal coordinate analysis (PCoA), also known as multidimensional scaling [35], to find a matrix *Z* such that the Euclidean distance between its *i*-th and *j*-th column is approximately equal to *D*(*i, j*). Therefore, the matrix *G* := *Z*^*′*^*Z* is of a Gram-type [36] as its entries are the Euclidean inner products between all the columns in *Z*.

Define *K*_*w*_ := *X*^*′*^Φ^*′*^ diag(*w*) Φ*X*; in particular, the entry associated with row-*x*_*i*_ and column-*x*_*j*_ of this matrix is precisely *k*_*w*_(*x*_*i*_, *x*_*j*_). Furthermore, *K*_*w*_ like *G* is also of a Gram-type because *K*_*w*_ = *Y* ^*′*^*Y*, with 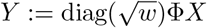. Ideally, we would like to select a weight vector *w* so that *K*_*w*_ = *G*; however, this is not generally possible. So instead, we pursue the next best option: *finding a vector w such that K*_*w*_ *approximates G as best as possible*, which we interpret as solving the optimization problem:

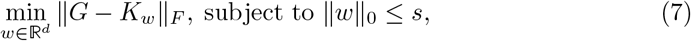

where ∥ · ∥_0_ is the pseudo-norm that counts the number of nonzero entries of the vector. The inclusion of the constraint ∥*w*∥ _0_ ≤ *s*, where *s* is a strictly positive user-defined integer, ensures that the optimization prioritizes sparse solutions, thereby enhancing the interpretability of the solution *w*. Figure 1 gives an overview of the approach we have just described.

**Fig 1.**
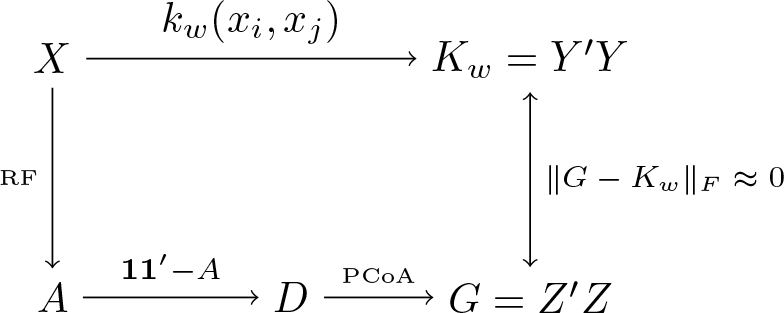
Illustration of the main mathematical objects and their relationships in our metric learning algorithm.

**Fig 2.**
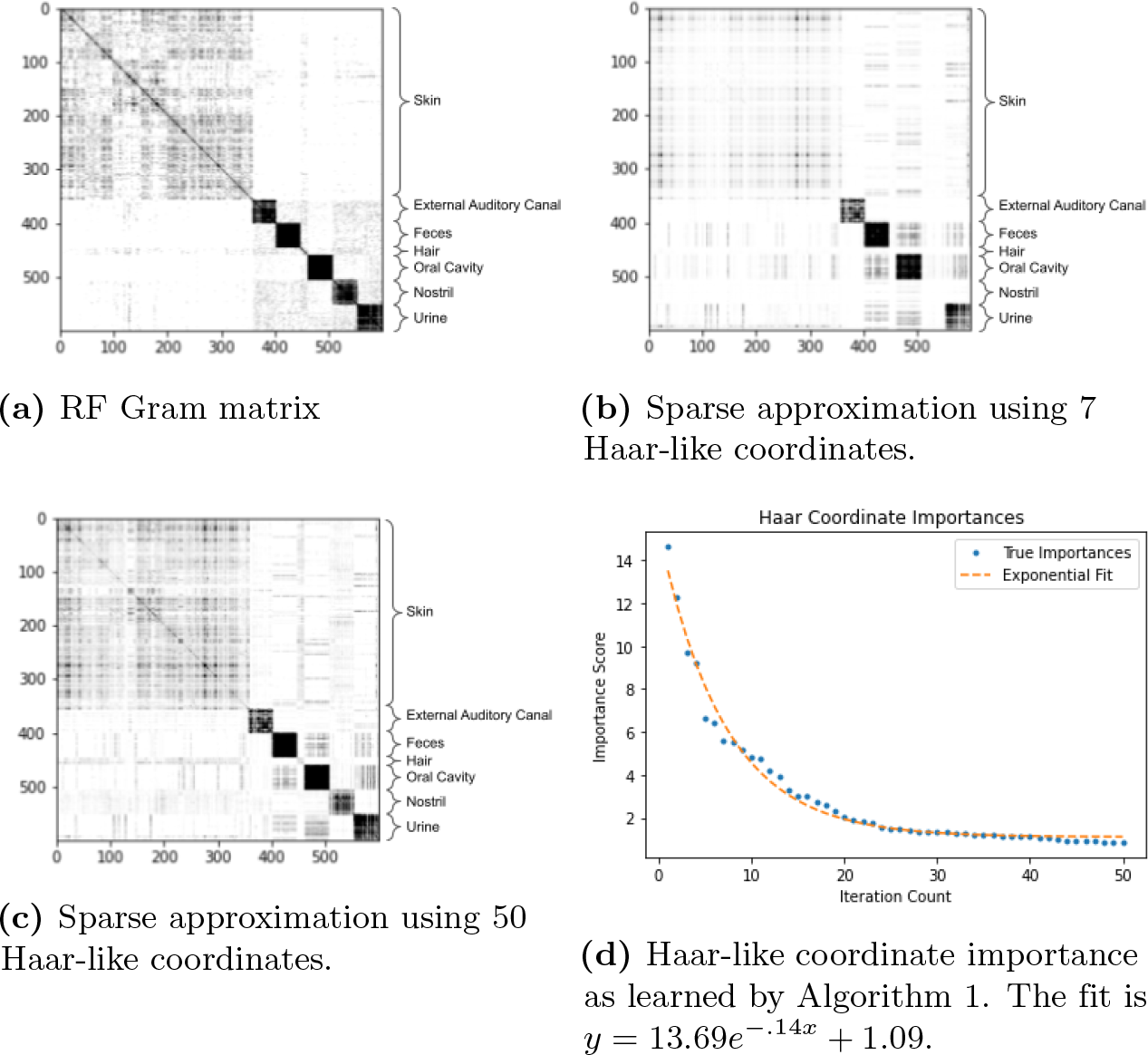
Sparse approximation of the RF Gram matrix from the Body Sites dataset.

It is worth noting that the identities, *G* = *Z*^*′*^*Z* and *K*_*w*_ = *Y* ^*′*^*Y*, prompt the approximation of Euclidean coordinates in *Z* by those in *Y*. Nevertheless, this alternative approach is unsuitable because it is not rotation-invariant, unlike the formulation based on Gram matrices in (7).

For any matrix *M*, let *M*_*i*_ denote its *i*-th column and vec(*M*) be the (column) vector obtained by stacking *M*_1_, *M*_2_, … up from left to right.

We can reformulate the optimization problem in (7) into a more computationally tractable one as follows. Since *K*_*w*_ is linear in *w*, there is a matrix 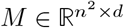 such that vec(*K*_*w*_) = *Mw*. In particular, since ∥ *G* −*K*_*w*_∥ _*F*_ = ∥ vec(*G*) − vec(*K*_*w*_) ∥ _2_, the minimization problem is equivalent to

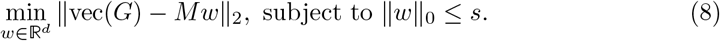

Further, the columns of *M* can be computed explicitly as follows. Let *e*_*i*_ denote the *i*-th vector in the canonical base of ℝ^*d*^; namely, *e*_*i*_(*j*) = ⟦*j* = *i* ⟧ for 1 ≤ *j* ≤ *d*. Then, we must have 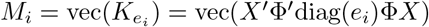. But diag(*e*_*i*_) is symmetric and idempotent; hence

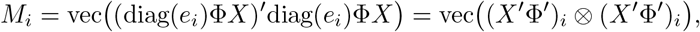

where ⊗ denotes the outer-product of vectors. Namely, *M*_*i*_ is the vectorization of the matrix obtained by the outer-product of the *i*-th row of Φ*X* with itself. We emphasize that Φ*X* is the matrix of Haar-like coordinates of the (unlabelled) data.

The optimization in equation (8) is a standard sparse approximation problem [37], where the matrix *M* is referred to as the “dictionary,” and the goal is to learn a sparse linear combination of the “dictionary elements” (i.e. columns of *M*) to best reconstruct a signal. In our setting, **in a dataset-specific manner**, the signal is the matrix *G*, which is a proxy for the discriminatory patterns learned by the RF, whereas *M* ‘s columns represent discriminatory patterns associated with the internal nodes of the reference phylogeny. The goal of the sparse approximation in (8) is therefore to **find the least number of Haar-like coordinates that can best explain the patterns learned by the RF**. We do this using Algorithm 1, a variant of the Matching Pursuit (MP) algorithm [37], in which the weights are constrained to be nonnegative.

#### Algorithm 1

**Non-negative Matching Pursuit Algorithm**

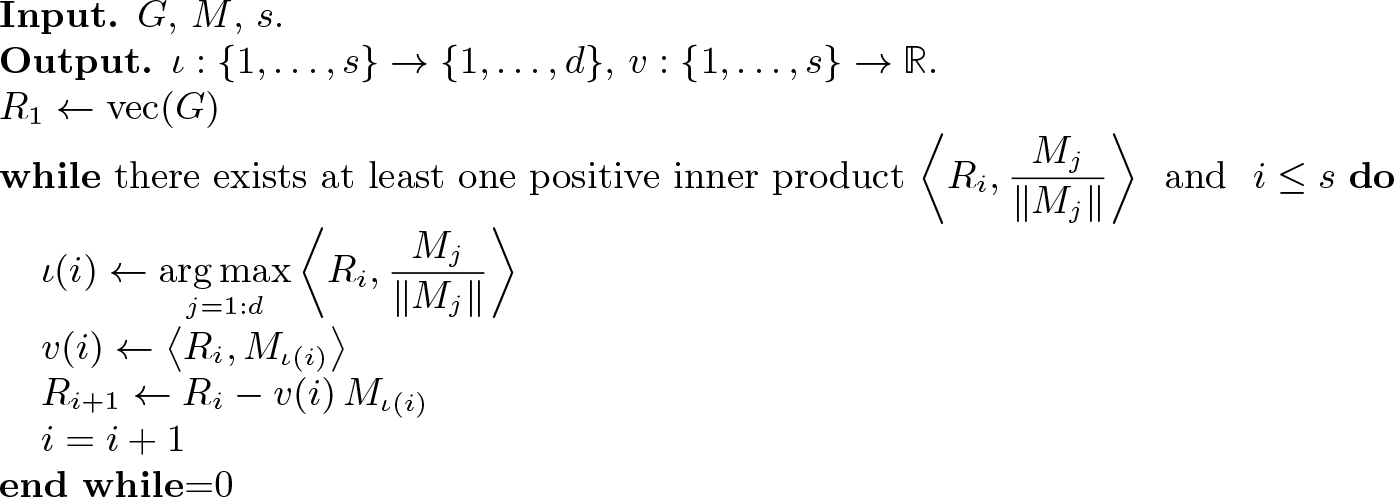

Algorithm 1 takes as input the signal *G*, the dictionary *M*, and a user-defined sparsification parameter *s*. It returns functions *ι* : {1, …, *s*} → {1, …, *d*} and *v* : {1, …, *s*} → ℝ such that the vector *w* with zero entries, except that *w*(*ι*(*i*)) = *v*_*i*_ for *i* = 1 : *s*, approximate solves the optimization problem in (8). The algorithm is greedy, which lets us choose exactly how sparse of a solution we want. Its key idea is to select iteratively the dictionary element with the largest projection along the signal, subtract this projection from the signal, and then repeat with the signal residual.

In principle, all inner products may be negative in the first iteration of the algorithm, in which case it will return no weights. Otherwise, as in traditional Matching Pursuit, the same column may be selected more than once. Nevertheless, we did not observe any of these anomalous behaviors when the sparsity was constrained to 10 or fewer coordinates.

The function *v* need not be decreasing, i.e. *v*(*i*) may be larger than *v*(*i* + 1). Nevertheless, the norm of *R*_*i*_ − *R*_*i*+1_ = *v*(*i*) *M*_*ι*(*i*)_ is a decreasing function of *i* (this follows from the non-constrained version of Matching Pursuit [38]) and a natural measure of the importance of the Haar-like coordinate with index *ι*(*i*); accordingly, we refer to | *v*(*i*) | · ∥ *M*_*ι*(*i*)_ ∥ as a **Haar-like coordinate importance**.

Because these learned weights are constrained to be nonnegative, the resulting Gram matrix, *K*_*w*_ = *X*^*′*^*W* ^2^*X* with 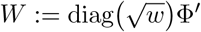 (definition 3), can be factored to find an associated Euclidean embedding with coordinates given by *WX*.

### The Haar-like Kernel as a Local Average Estimator

The question remains: **how consistent is the adaptive Haar-like surrogate model with the original random forest?**

In this section, we detail how the adaptive Haar-like kernel can be used as a local average estimator, similar to the KeRFE, to obtain estimates of unlabelled data points. Later, this allows us to benchmark our metric against the original random forest and another interpretable model, CoDaCoRe [39].

Again consider *n* labeled samples (*x*_*i*_, *y*_*i*_) ∈ (ℝ^*d*^, ℝ) with {*x*_*i*_}_*i*=1:*n*_ collected into a data matrix *X* ∈ ℝ^*d×n*^. We train the random forest using these labeled samples. Suppose we are then given *m* new unlabelled samples 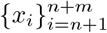 that are appended to the data matrix to form 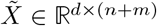. First, we construct estimates for these new points using the trained random forest. We then recover the random forest affinities between **all** points to construct the full affinity matrix *Ã*_*RF*_ ∈ℝ^(*n*+*m*)*×*(*n*+*m*)^. (Recall that affinity matrices are symmetric.) Next, we apply the metric learning algorithm to this affinity matrix to recover Gram matrix *K*_*w*_ and associated Haar-like coordinates 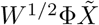. The Euclidean distances between these points are computed to form a Euclidean distance matrix *D*_Haar_. Values in this matrix are threshold to a maximum of one, allowing us to form the Haar Affinity: *A*_Haar_ = 1 − *D*_Haar_. Using this learned Haar Affinity, the surrogate estimate for the RF is:

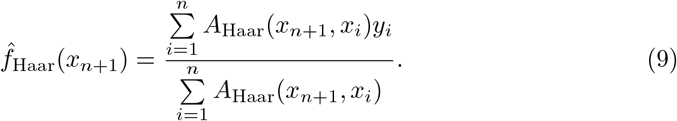

By replacing the RF affinity with our Haar-like affinity, we now have constructed an **interpretable** surrogate for the RF estimator: *estimates are made by comparing to neighbors and neighbors are determined by comparing the learned Haar-like coordinates*. Ahead, we demonstrate that this surrogate has comparable performance to the original RF.

## Results

In this section, our goals are twofold: to demonstrate the use of the adaptive Haar-like distance as an exploratory tool in metagenomics datasets (**Model Demonstration**), and to verify that our model is a suitable approximation of the original random forest (**Model Validation**).

For the model demonstration, we apply the adaptive Haar-like distance to four datasets spanning categorical and continuous labels across varied biological settings. We examine the learned Haar-like coordinates for each dataset, producing visualizations and associating them with known biological contexts. Next, for the validation, we benchmark the adaptive Haar-like distance classification performance against the standard RF and another interpretable classifier, CoDaCoRe [39].

Our analyses are based on 16s rRNA sequence data; however, our methodology applies to whole genome sequence (WGS) and other sequence types as long as a reference phylogeny is provided. We used Greengenes 97% as the reference phylogenetic tree and **we report Haar-like coordinates by their associated index in a post-order traversal of the tree starting with index 0**.

Due to the correspondence between the Haar-like basis and the set of internal nodes in the reference phylogenetic tree, we can construct a helpful visualization for the learned metric over a dataset. We display the **phylogenetic spectrogram** associated with our metric by shading clades according to their learned weights (see Definition 1.5 and Figures 3,9,14, and 19). These figures were generated using iTOL [40].

**Fig 3.**
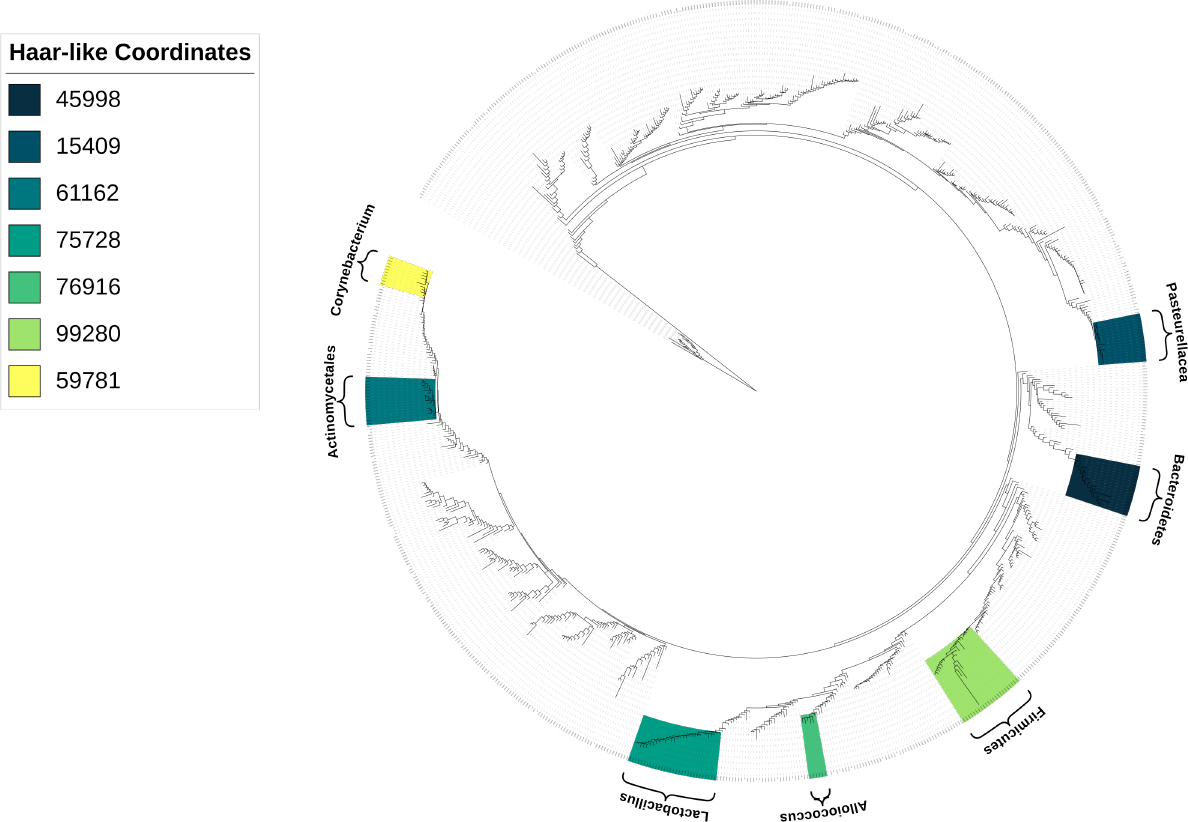
The seven most important Haar-like coordinates of the Body Sites dataset visualized on Greengenes 97%.

Our general methodology has two user-controlled tuning parameters: the sparsity *s* (i.e., the number of Haar-like coordinates to recover), and the minimum RF bin size (i.e., the minimum number of samples for a node to be considered a leaf in the original RF), initially set during the training of the original RF. In the classification setting, we always set the minimum RF bin size to 1 for optimal performance. However, in the regression setting, too small a bin size may reduce the RF affinity between similar samples and can make it more difficult for our algorithm to recover clustering patterns. Accordingly, we set the bin size at 10% of the total sample count, noting that optimal bin size may require further experimentation.

### Model Demonstration

The purpose of this section is to introduce the adaptive Haar-like distance as an exploratory tool to **link differences in environmental characteristics (given by sample labels) to variations in clade abundances**. We show that across a diverse range of microbial environments, our metric produces embeddings that display strong clustering (in the classification setting) and strong gradients (in the continuous setting) with respect to these sample labels.

A particularly useful aspect of our metric is that, due to Defition 1.5, the Haar-like coordinates can be treated as Euclidean ones. These coordinates correspond to speciation events in the reference phylogenetic tree, facilitating direct visualization of the relationship between changes in clade abundances and directions within the embedding through a biplot [41], where loadings are associated with Haar-like coordinates. Notably, by constraining the number of Haar-like coordinates, we achieve a distinct advantage over existing phylogenetic *β*-diversity metrics: the resulting embedding can be explained **exactly** by a small number of clades.

We underscore that our aim here is not to replicate a full scientific analysis of these datasets but to demonstrate that our metric recovers Haar-like coordinates associated with biologically significant clades. With this in mind, for each dataset, we select ahead of time a sparsity parameter *s* and link the top *s* Haar-like coordinates to established taxonomical annotations (given by the Greengenes 97% taxonomy) by identifying the lowest taxonomic classification that encompasses all members of the corresponding clade. While further Haar-like coordinates may be relevant—depending on the dataset—we limit our discussion to these top *s* coordinates. Nevertheless, an in-depth analysis of the relevant Haar-like coordinates should pay close attention to the comparison of abundances of the left and right descendants of the corresponding internal nodes. For instance, an observed increase in a Haar-like coordinate does not necessarily imply increased abundances of all its descendants. Instead, recall from Theorem 1.1 that the Haar-like coordinate values represent the difference between the abundances of left and right subtrees.

### Dataset 1: Body Sites

The first dataset we analyze comes from “Bacterial community variation in human body habitats across space and time” [42]. This study “surveyed bacteria from up to 27 sites in seven to nine healthy adults on four occasions” resulting in a total of 600 samples. For our analysis, we grouped these into 7 primary body habitats: skin, external auditory canal, feces, hair, oral cavity, nostril, and urine.

Training the RF on all 600 labeled data points, we form the RF Gram matrix *G* shown in Figure 2a. Here, the indices have been sorted by body habitat. For this dataset, motivated by the fact that there are seven body habitats, we first applied the non-negative Matching Pursuit algorithm with *s* = 7 to recover the seven most important Haar-like coordinates. The associated Gram matrix *K*_*w*_ constructed from these coordinates is shown in Figure 2b and the associated phylogenetic spectrogram in Figure 3. We also display the Gram matrix resulting from the first 50 coordinates in Figure 2c. In both cases, we find a good reconstruction of the true RF affinity with only a small amount of additional noise. Figure 2d displays the importance of these top 50 Haar-like coordinates. We note that the exponential decay of these importances implies low dimensional embeddability of the data and indicates the efficiency of our adaptation of Matching Pursuit (Algorithm 1) in choosing relevant Haar-like coordinates.

For illustration, Figure 4 displays how the first seven Haar-like coordinates combine to capture the vast majority of the classification pattern (excluding hair samples) seen in the RF Gram matrix (Figure 2b). We note that hair samples make up only about ∼ 2% of the dataset, so their lack of distinction with only seven Haar-like coordinates is not surprising.

**Fig 4.**
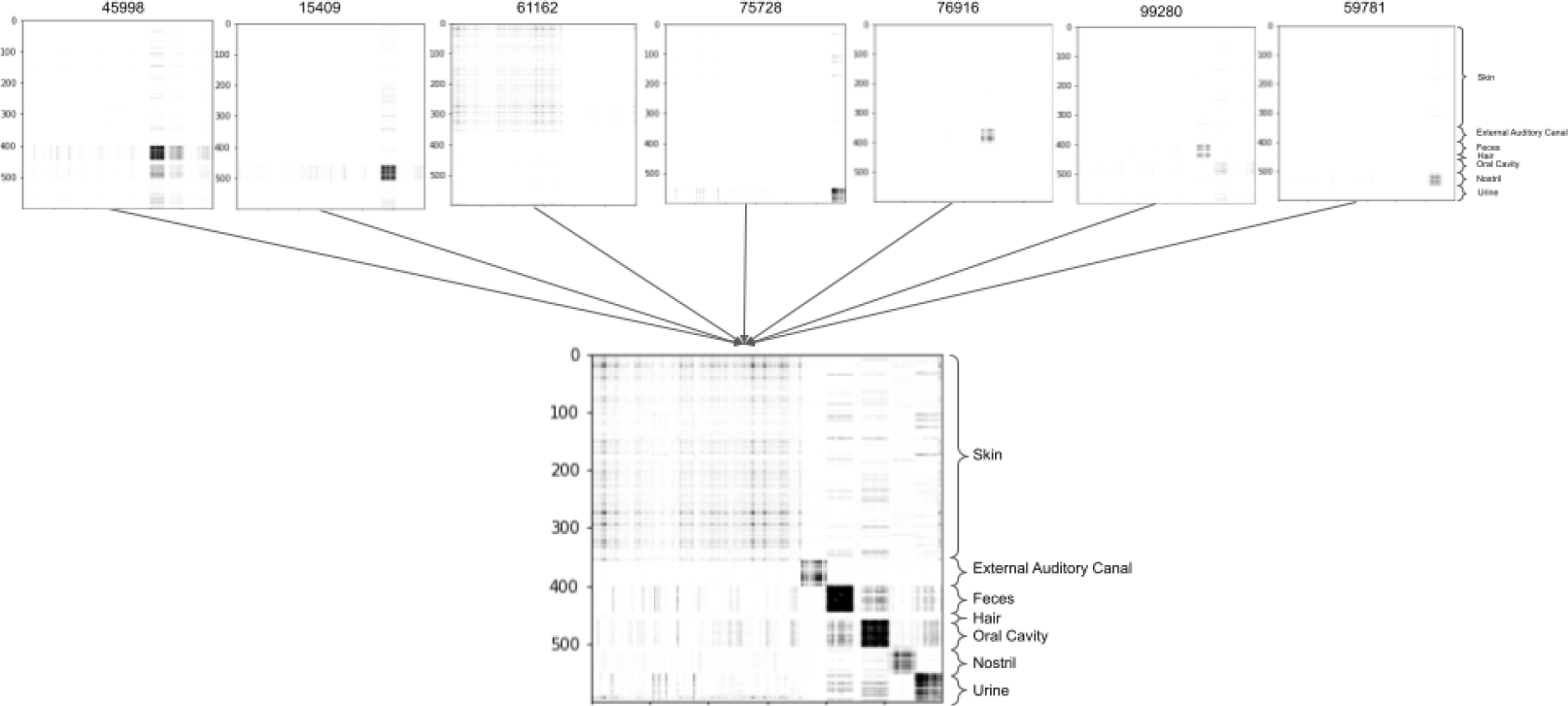
Reconstructed RF Gram matrix of the Body Sites dataset using the seven most dominant Haar-coordinates. These have indexes 45998, 15409, 61162, 75728, 76916, 99280, and 59781 in a post-order traversal of Greengenes 97%.

To confirm that our algorithm is recovering biologically meaningful splits in Greengenes 97%, we further examine these first seven selected Haar-like coordinates to assess their relevance to the habitat of interest. To aid in our analysis, Figure 5 displays boxplots of these seven coordinates in the different body habitats.

**Fig 5.**
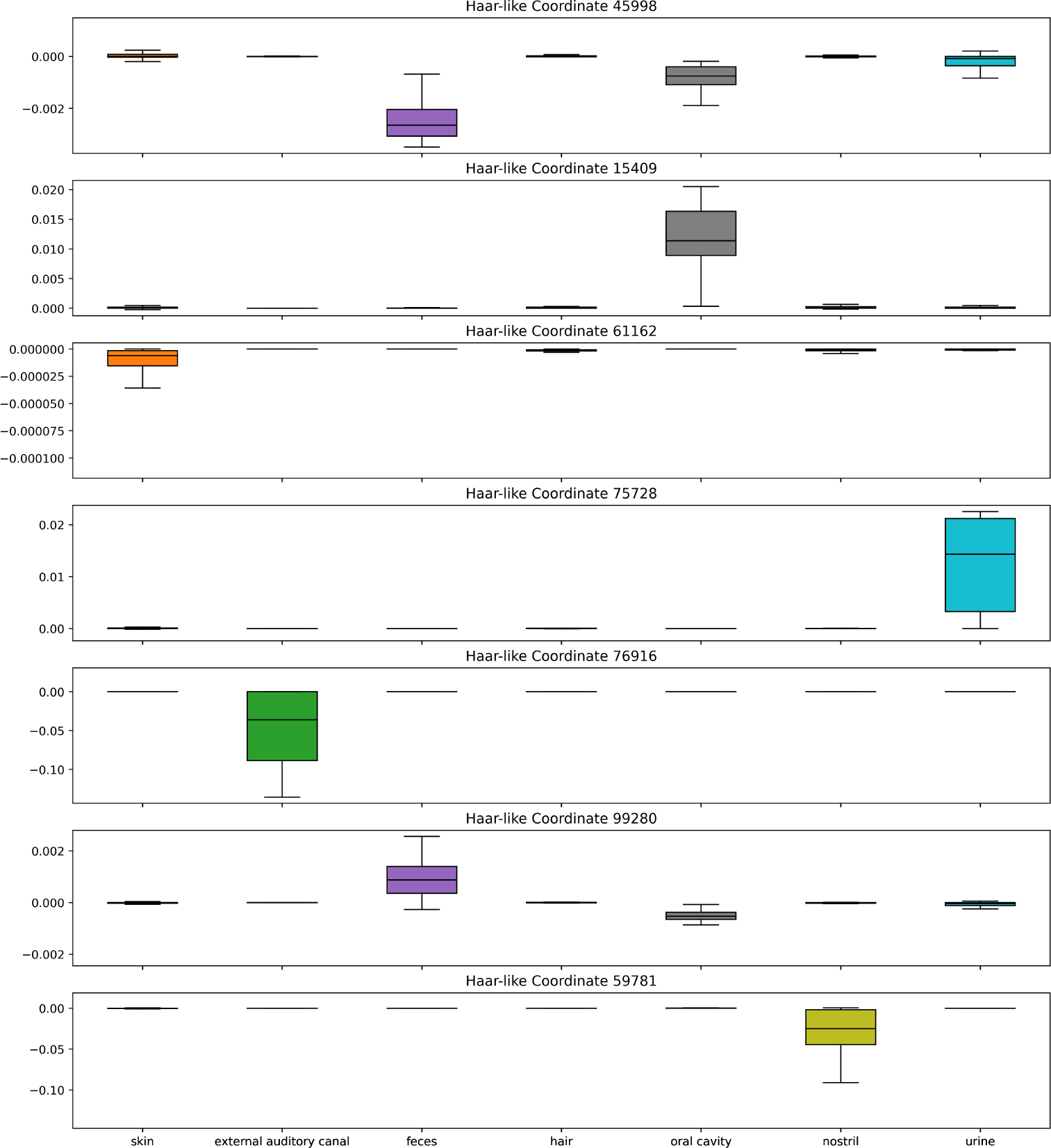
Box plots of the top seven Haar-like coordinates across the Body Sites dataset.

The dominant Haar-like coordinate (45998) strongly localizes fecal samples (with a negative coordinate value) and also corresponds, to a lesser extent, with oral cavity and urine samples. Notably, the descendants of node 45998 are classified as Bacteroidetes, a phylum well known to be found in the human gut, but also in the mouth and urine [43].

The second selected Haar-like coordinate (15409) localizes oral cavity samples. On examination of the phylogeny, this clade consists entirely of the Pasteurellaceae family, which has been identified in human supragingival plaque samples [44].

The third selected Haar-like coordinate (61162) corresponds to the order Actinomycetales. As seen in Figure 5, this coordinate strongly localizes the skin samples. The literature supports that Actinomycetales, specifically the order genus Actinomyces, appear in the skin [45].

The fourth selected Haar-like coordinate (75728) localizes urine samples and consists entirely of the genus Lactobacillus, which has been found in both male and female urine [46, 47].

The fifth selected Haar-like coordinate (76916) localizes the external auditory canal and consists entirely of the genus Alloiococcus, which has been established as part of the typical outer ear microbiome [48].

The sixth selected Haar-like coordinate (99280) again localizes fecal and oral cavity samples. This clade contains the Firmicutes phylum, which are well-known members of the gut and oral microbiome [49].

Finally, the seventh selected Haar-like coordinate (59781) strongly localizes nostril samples. This clade consists entirely of the Corynebacterium genus, which is known to be a dominant bacteria of the nose [50].

Based upon these top seven Haar-like coordinates, we then apply principal component analysis (PCA) to reduce to three dimensions for visualization. Comparing our PCA embedding (Figure 6d) to the PCoA embeddings associated with unweighted UniFrac, weighted UniFrac, and the Haar-like distance (Figures 6a,6b,6c) we see that our metric obtains better clustering by bodysite. Because this dataset has a large number of classes, it can be difficult to see all of the class separations in the biplot. For this reason, we also display the **normalized** PCA embedding, resulting from a rescaling of the Haar-like coordinates, in Figure 7. In the normalized biplot, we can see all seven loadings, and it is clear that our metric is recovering Haar-like coordinates that align well (either in the positive or negative direction) with different classes in the embedding. For example, coordinate 15409, which was linked to Supragingival plaque, points exactly in the direction of significant variation for the oral cavity.

**Fig 6.**
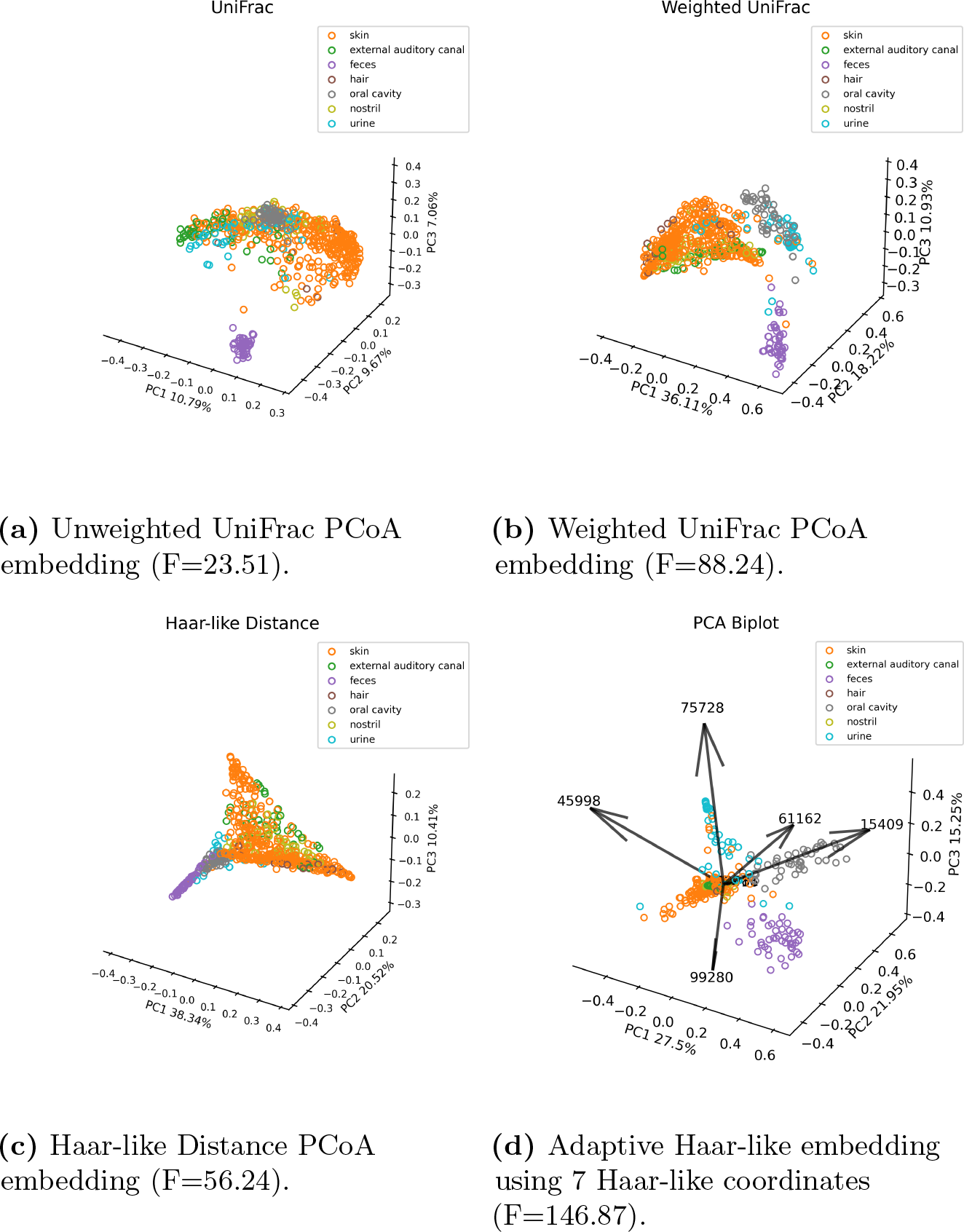
Comparison of the Adaptive Haar-like embedding to various phylogenetic *β*-diversity metrics in the Body Sites dataset. Pseudo F-statistics are reported to quantify clustering.

**Fig 7.**
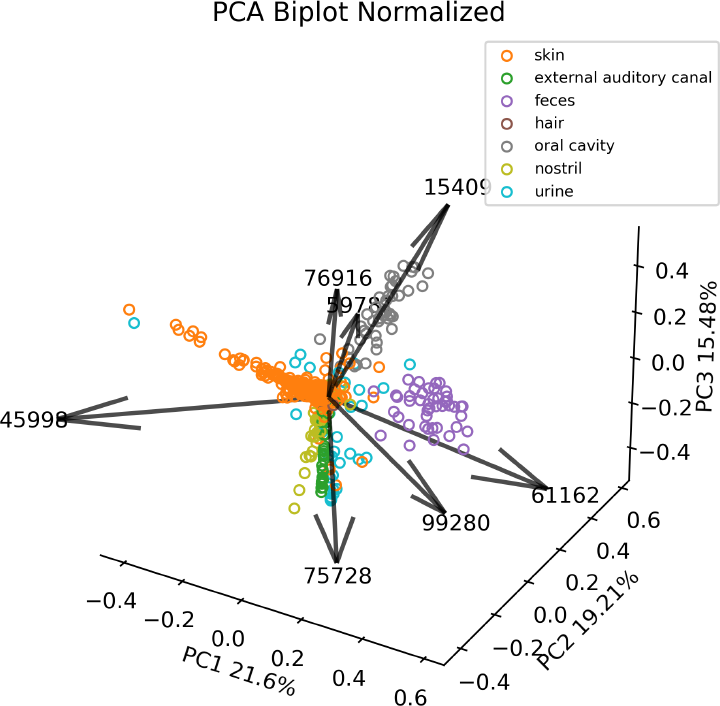
Normalized PCA of adaptive Haar-like distance using seven coordinates in the Body Sites dataset.

We can quantify how well each metric clusters the body habitats by the PERMANOVA pseudo-F test statistic [51] applied to the corresponding distance matrices, which is a measure of clustering strength estimated by comparing within-group variability to between-group variability; a higher value indicates stronger clustering. As expected, the embedding associated with the adaptive Haar-like distance has the highest test statistic and, by this measure, recovers the best clustering of body habitats among the various phylogenetic *β*-diversity metrics.

Altogether, these seven coordinates are enough to cluster all body habitats except for the hair samples, which, as we have mentioned, represent a too small fraction of the dataset for localization with only seven Haar-like coordinates. Though separating body habitats may be a relatively trivial classification task, our method recovers coordinates that align almost perfectly with the various body habitats and outperforms existing metrics using just a few coordinates.

Next, we show that our algorithm maintains strong performance even on classification tasks deemed far more challenging by current metagenomic analyses.

### Dataset 2: Autism

We turn our attention to the dataset from “Altered gut microbial profile is associated with abnormal metabolism activity of Autism Spectrum Disorder” [52]. This study compared fecal samples from 143 individuals diagnosed with autism spectrum disorder (ASD) against 143 control subjects, matched for age and gender.

In Figure 8, we display the sparse approximation of the RF Gram matrix using either 3 or 50 Haar-like coordinates. We observe clustering among ASD and control subjects, but the reconstruction displays more noise than the previous dataset. This should be expected because the Body Sites data (Dataset 1) compared samples from distinct body habitats, which are known to harbor different microbial communities [42], while Dataset 2 contains samples from the same habitat (feces). Consequently, it may be harder to find features that strongly distinguish the two groups.

**Fig 8.**
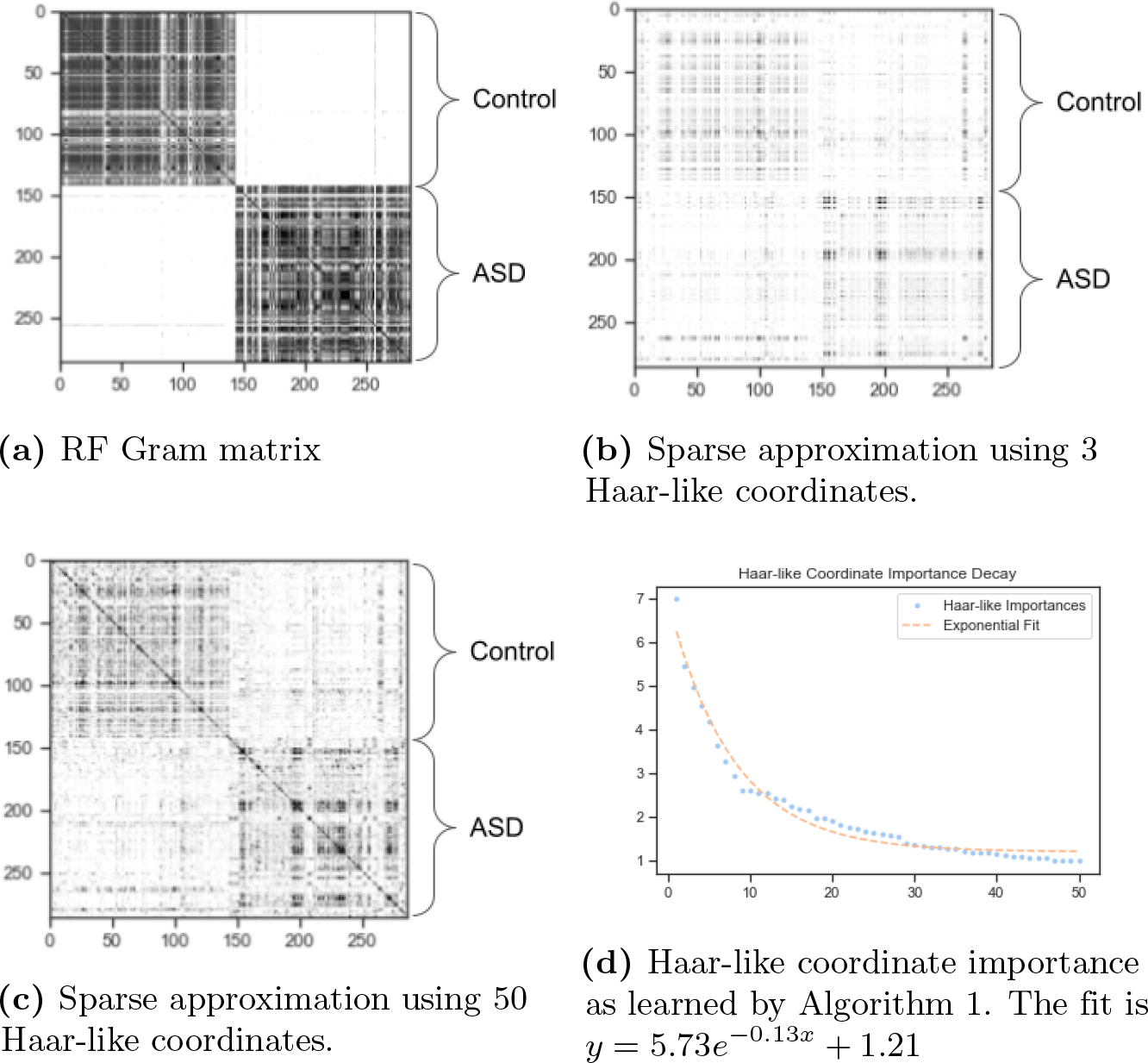
Sparse approximation of the RF Gram matrix from the Autism dataset.

**Fig 9.**
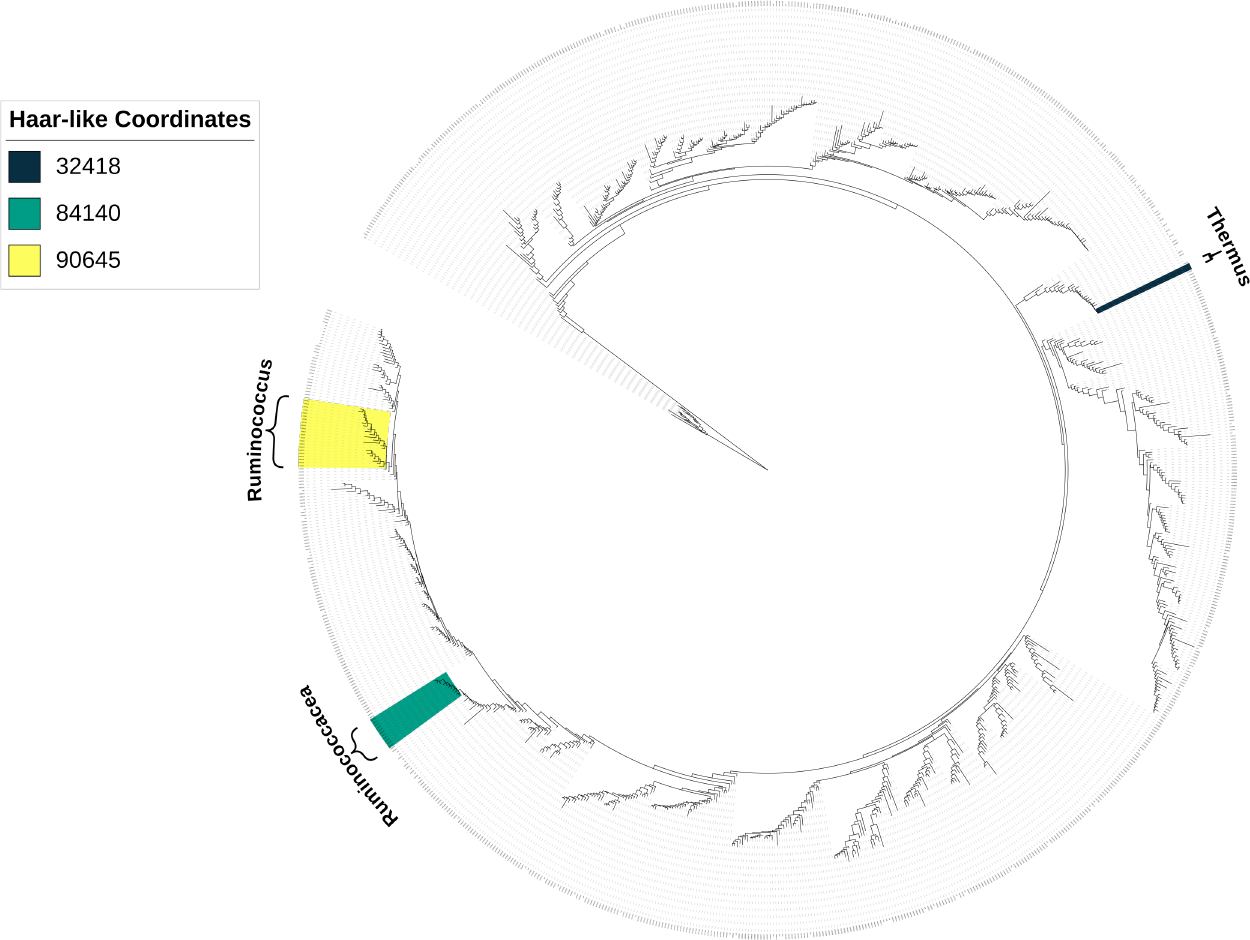
The three most important Haar-like coordinates of the Autism dataset visualized on Greengenes 97%.

As seen in Figure 12, the dominant coordinate (32418) strongly distinguishes the ASD patients. This clade consists entirely of the genus Thermus, which has been observed to differ significantly in ASD patients compared to controls [53]. For the next prominent coordinate (84140), the associated clade is made up of an unclassified genus within the Ruminococcaceae family. Finally, for the third coordinate (90645), every member of the corresponding clade is classified as Ruminococcus, a genus that has been associated with ASD in the original study of this dataset [52], as well as in other research [54–56].

In the phylogenetic spectrogram (Figure 9), we note that the second and third coordinates (84140 and 90645) are more closely related (both belonging to the order Clostridiales) than the dominant coordinate (32418), which descends from Deinococci, a class of extremophiles. The role of these extremophiles, such as Thermus, in Autism spectrum disorder is not well understood, yet our methodology highlights their potential significance in this context.

For this dataset, we only compute the embedding associated with the three most dominant Haar-like coordinates. While we do not achieve the same quality of class separation as the previous dataset, the adaptive metric still achieves the best clustering among the tested metrics as indicated by the PERMANOVA statistics (Figure 10). This improvement in clustering compared to the (non-adaptive) Haar-like distance serves as compelling evidence for the role of weight optimization over the Haar-like coordinates to capture relevant differences in microbial composition.

**Fig 10.**
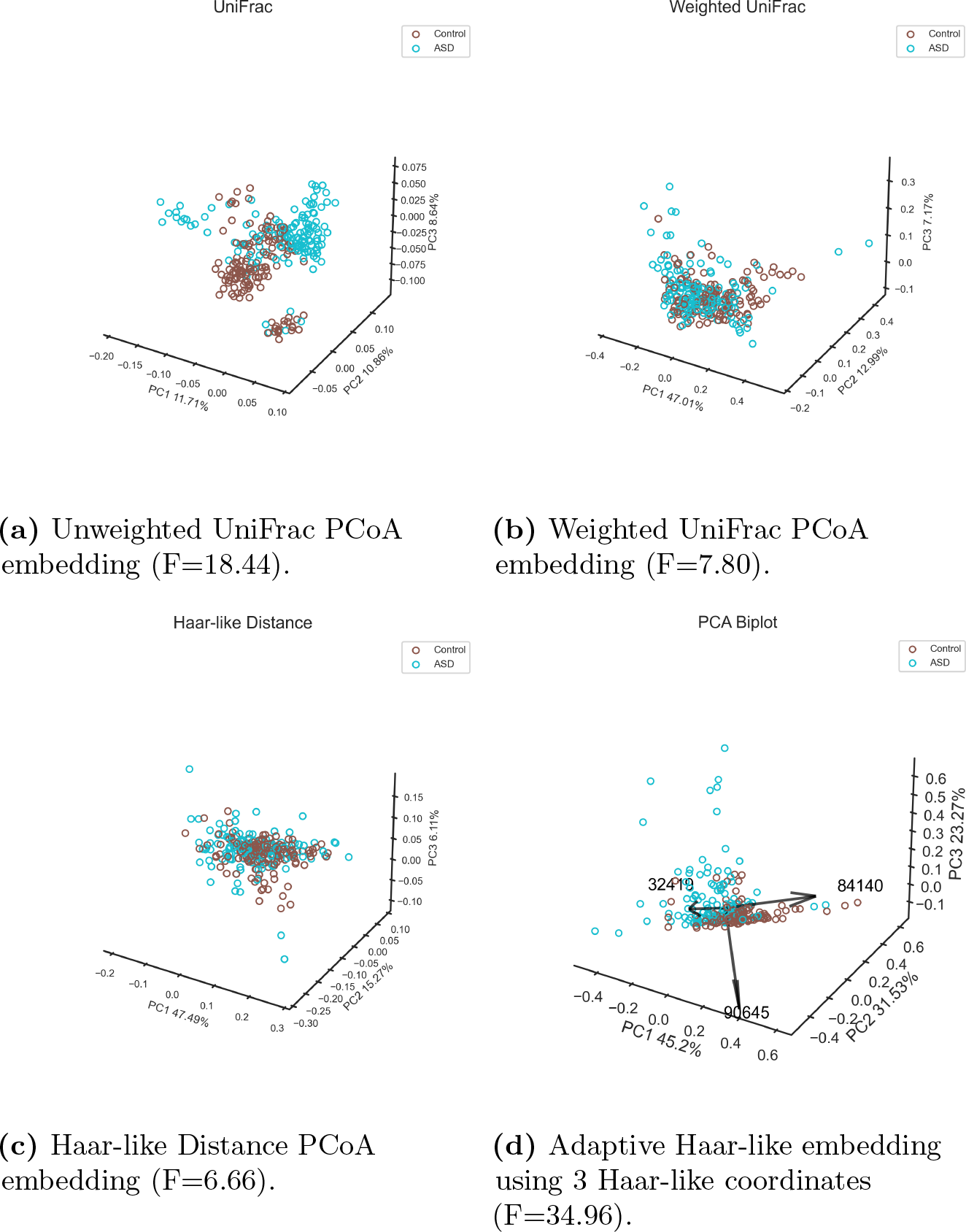
Comparison of the Adaptive Haar-like embedding to both variants of UniFrac and the Haar-like distance in the Autism dataset. Pseudo F-statistics are reported to quantify clustering.

Similarly to the previous dataset, we display the normalized PCA biplot in Figure 11 where the class separation is more visible. In both this plot and the non-normalized PCA, we see that coordinates 90645 and 32418 are nearly orthogonal: 90645 captures the variation in ASD patients, while 32148 corresponds to control patients.

**Fig 11.**
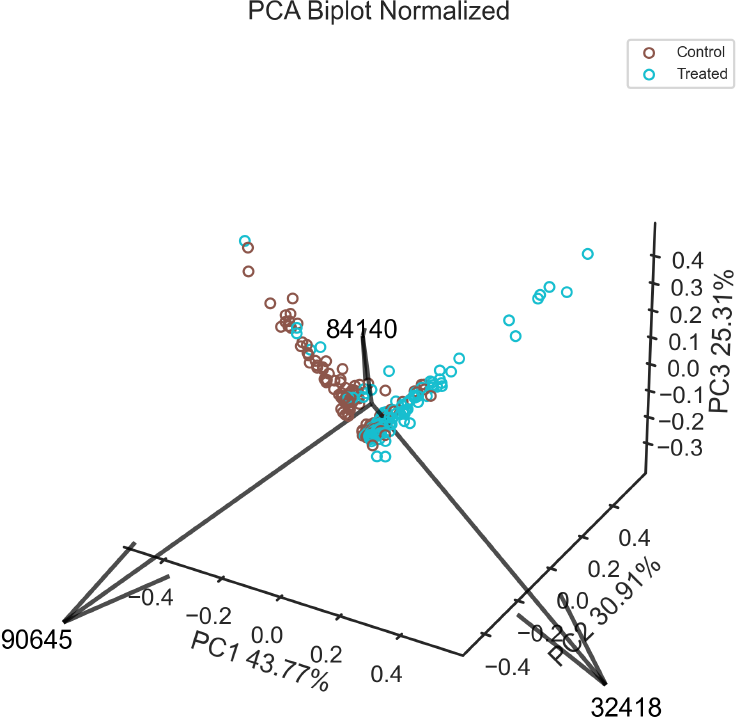
Normalized PCA of adaptive Haar-like distance using three coordinates in the Autism dataset.

**Fig 12.**
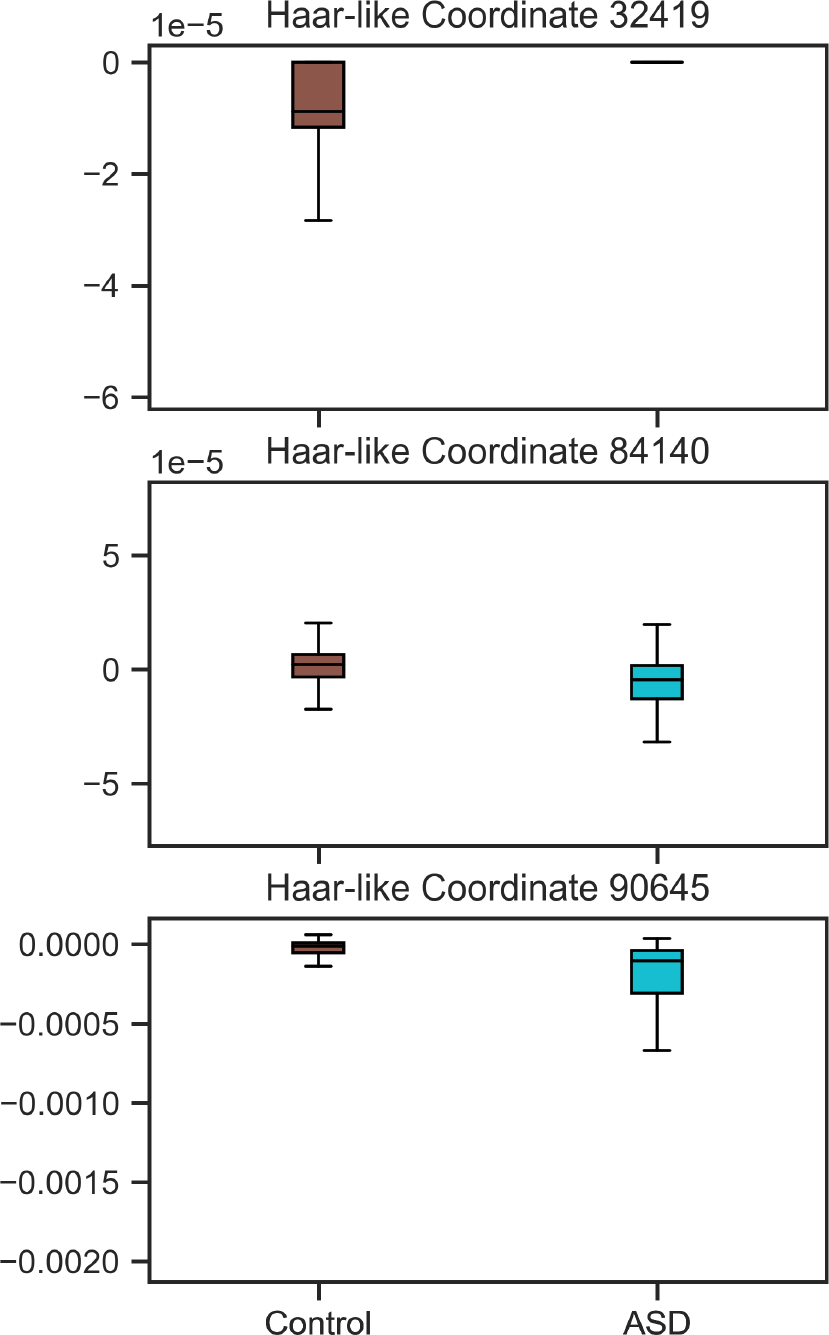
Boxplot of the Dominant 3 Haar-like coordinates in the Autism dataset.

Next, we demonstrate our methodology in a regression setting.

### Dataset 3: Deepwater Horizon Oil Spill

The first dataset with continuous labels that we consider “investigated the impact of oil deposition on microbial communities in surface sediments collected at 64 sites” affected by “the Deepwater Horizon oil spill in the spring of 2010” [57]. For our analysis, we consider each sample’s distance from the wellhead.

As seen in Figure 13a, when computing the RF Gram matrix, we find high inner product values, corresponding to high similarity, along a wide diagonal band with some faint off-diagonal noise. Interestingly, as seen in Figures 13b-c, when approximating the RF Gram matrix with either 4 or 50 Haar-like coordinates, we recover two clusters along the diagonal that are faintly present in the RF Gram matrix, but we do not recover as much of the diagonal band structure. Nevertheless, as seen in Figure 16d, the adaptive Haar-like distance still displays a very clear gradient with respect to sample distance.

**Fig 13.**
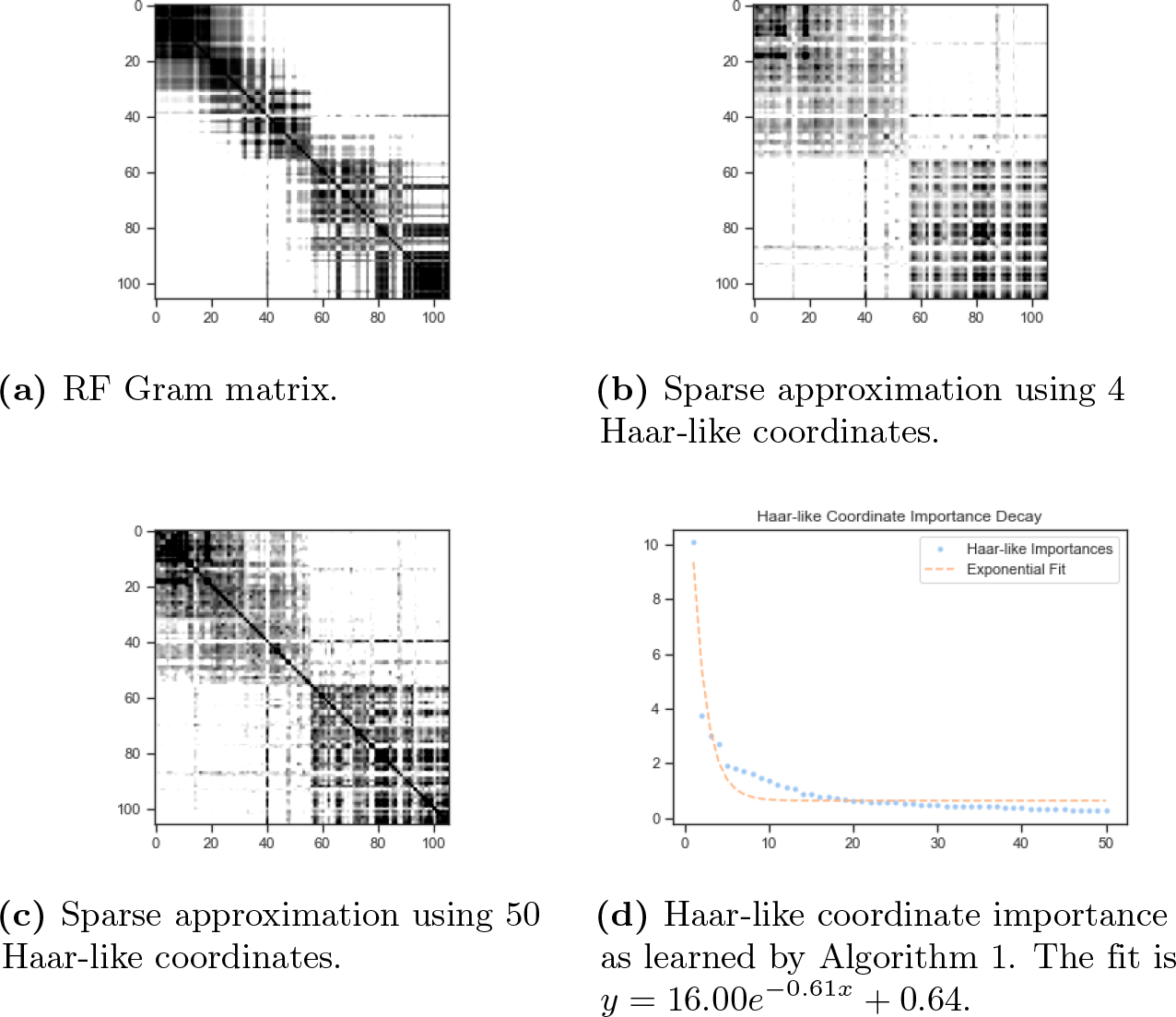
Sparse approximation of the RF Gram matrix from the Deepwater Horizon oil spill dataset.

**Fig 14.**
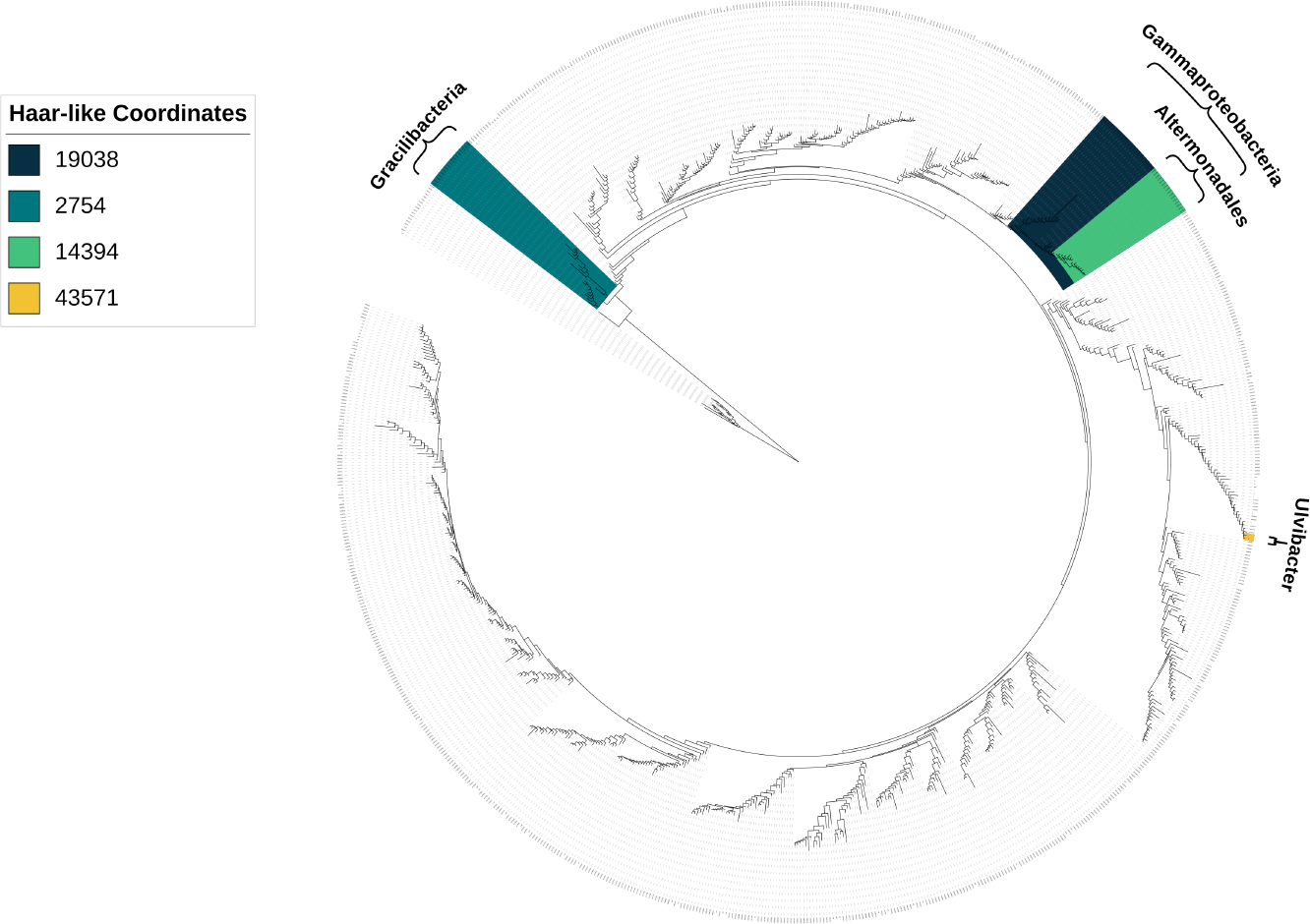
The four most important Haar-like coordinates of the Deepwater Horizon oil spill dataset visualized on Greengenes 97%.

**Fig 15.**
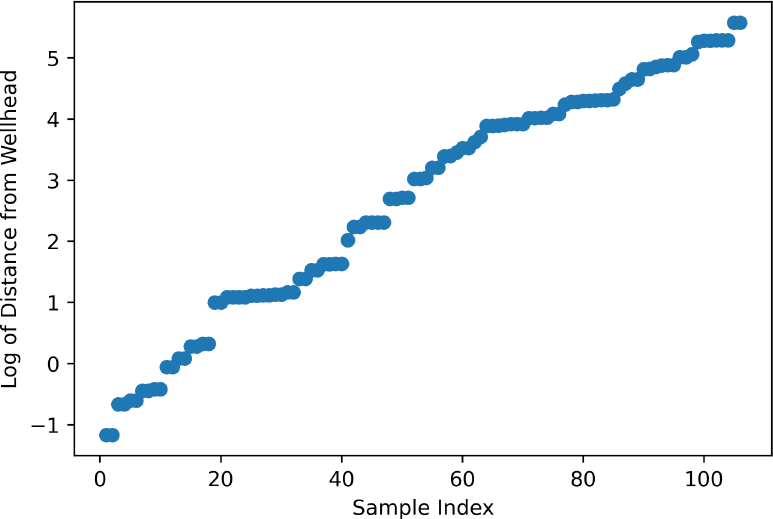
Logarithm of sample distances from the wellhead in the Deepwater Horizon oil spill dataset.

**Fig 16.**
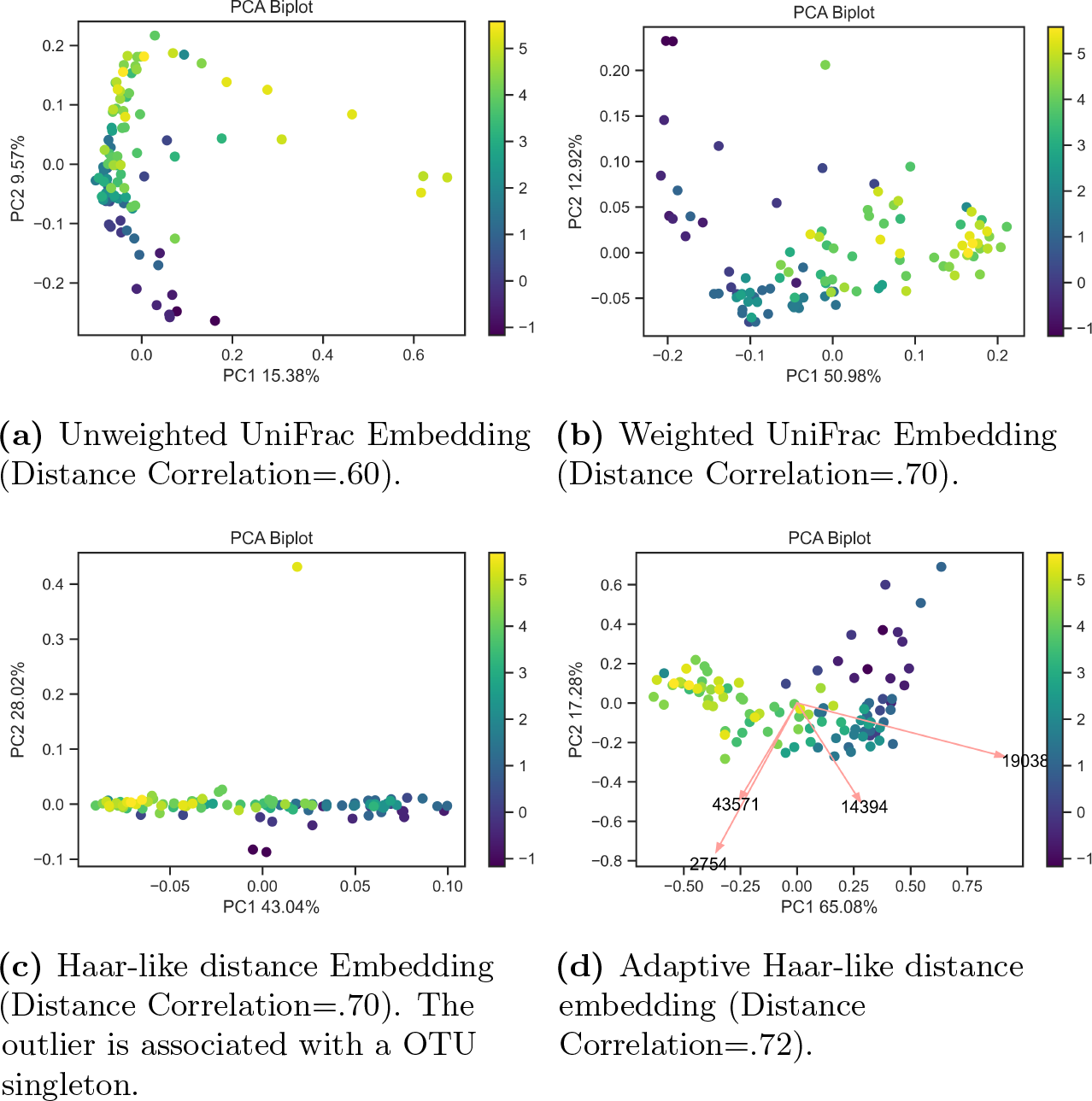
Comparison of the Adaptive Haar-like embedding to unweighted and weighted UniFrac in the Deepwater Horizon oil spill dataset.

In this dataset, the dominant Haar coordinate (19038) consists entirely of the class Gammaproteobacteria. As seen in Figure 17, this Haar-like coordinate decreases with distance from the contamination site, and as seen in Figure 16d, the corresponding loading aligns well with the distance gradient. This is consistent with the observation in the original publication [57] that an uncultured Gammaproteobacterium OTU and a Colwellia taxon had high relative abundances in highly contaminated samples, but low relative abundances elsewhere. The second Haar-like coordinate (2754) consists entirely of the phylum Gracilibacteria, which has been identified and examined in the context of oil spills previously [58]. The third Haar-like coordinate (14394) consists of the Alteromonadales order, which, as seen in Figure 14, descends from the clade corresponding to the dominant coordinate (19038). Altermonadales abundance has also been identified and examined in oil-contaminated samples in [59]. Finally, the fourth Haar-like coordinate (43571) consists entirely of the genus Ulvibacter, which is known to be a hydrocarbon-degrading bacteria [60].

**Fig 17.**
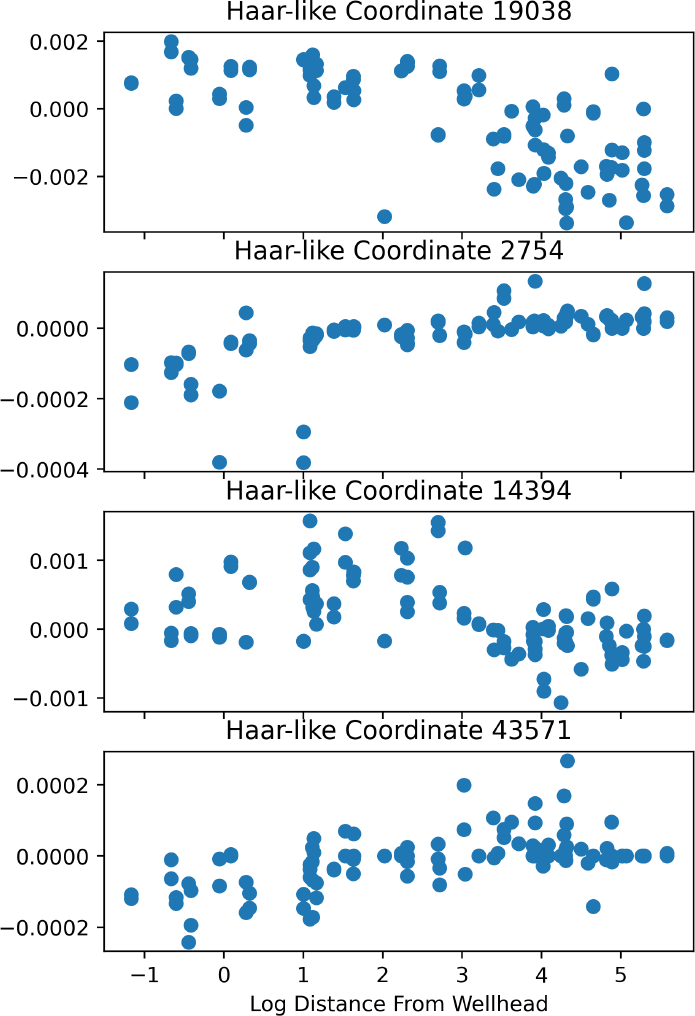
Plots of the top four Haar-like coordinates across samples of the Deepwater Horizon oil spill dataset.

These learned Haar-like coordinates all correspond to clades that play a role in oil degradation, and as we show next, together, they are sufficient to accurately cluster samples based on their distance from the wellhead.

Constructing the Adaptive Haar-like embedding using the top four Haar-like coordinates, we see a very clear gradient with respect to distance. For both the biplot (Figure 16d) and the plots of the individual Haar-like coordinates (Figure 17), we take the logarithm of the sample distances in order to approximately linearize the distances (see Figure 15). This has no effect on the resulting analysis and serves only for gradient visualization with respect to sample distance (to the wellhead) on a linear scale.

To quantify the strength of the gradients in the regression setting, we use distance correlation [61]. As opposed to the traditional notion of correlation, distance correlation captures nonlinear associations, and a distance correlation of zero is equivalent to probabilistic independence. Comparing the distance correlation between the true wellhead distances and the various phylogenetic *β*-diversity metrics, we find, as reported in Figure 16, that the adaptive Haar-like distance has the highest distance correlation among the four *β*-diversity phylogenetic metrics.

Next, we demonstrate our metric on samples taken from increasing depths in a microbial mat.

### Dataset 4: Microbial Mat

The final dataset we examine is from “Trade-offs between microbiome diversity and productivity in a stratified microbial mat” [62]. This study “evaluated the spatial relationships of productivity and microbiome diversity in a laboratory-cultivated photosynthetic mat.” For our analysis, we consider microbial abundances at different depths (1.5mm to 17.5mm) within the mat.

First, comparing the reconstructed matrices using either two or fifty Haar-like coordinates (Figure 18a-c), we see that two coordinates are enough to recover the outer ends of the diagonal band in the RF Gram matrix, corresponding to shallow or deep samples, but misses some similarity present among intermediate depths. However, just these top two coordinates are enough to recover a very clear gradient with respect to depth, as seen in Figure 21d. In fact, even though the weighted UniFrac embedding produces a strong gradient, the adaptive Haar-like distance resulting from these two coordinates achieves an even higher distance correlation with the true sample depths as seen in Figure 18d).

**Fig 18.**
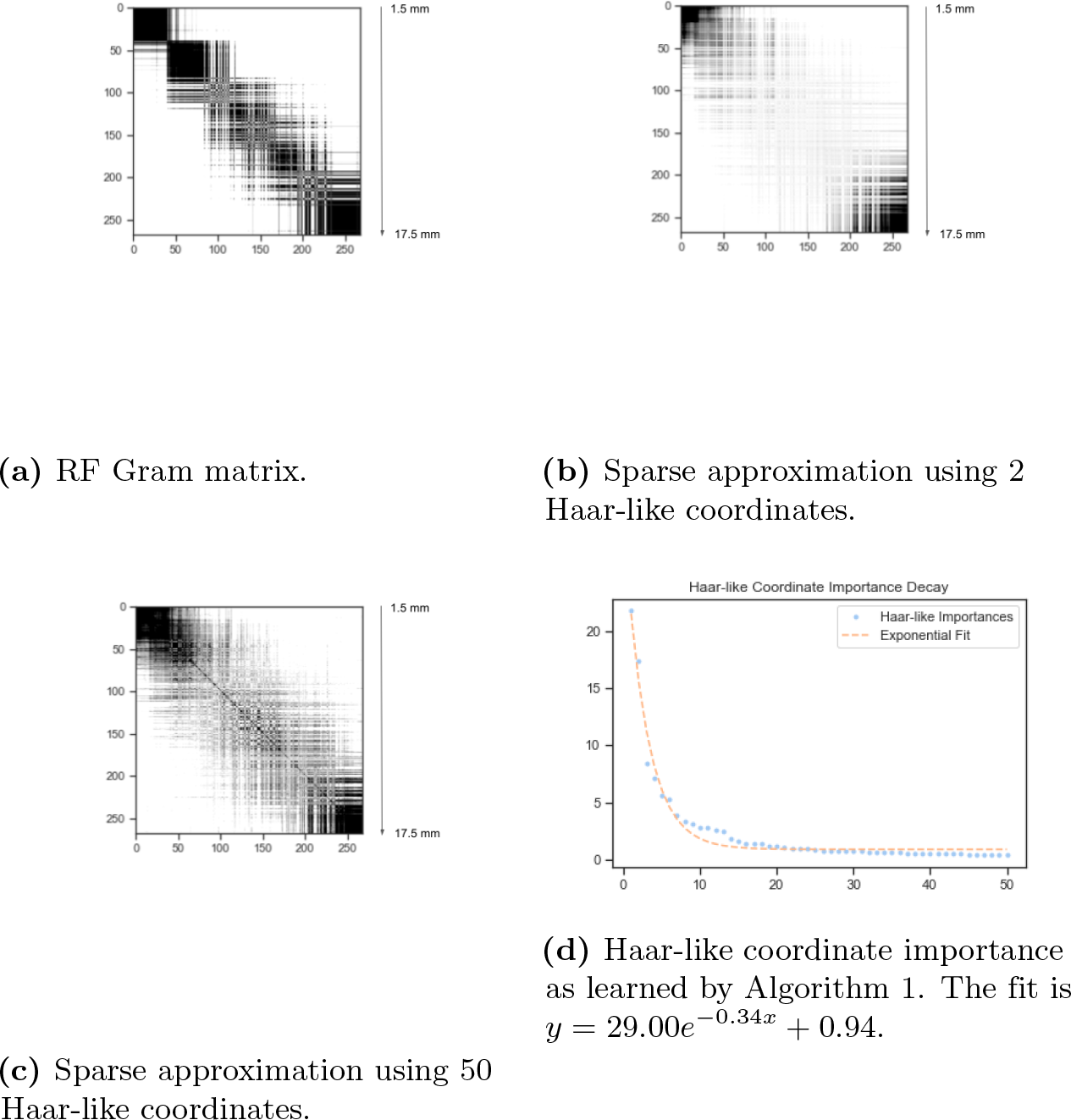
Sparse approximation of the RF Gram matrix from the Microbial Mat dataset.

**Fig 19.**
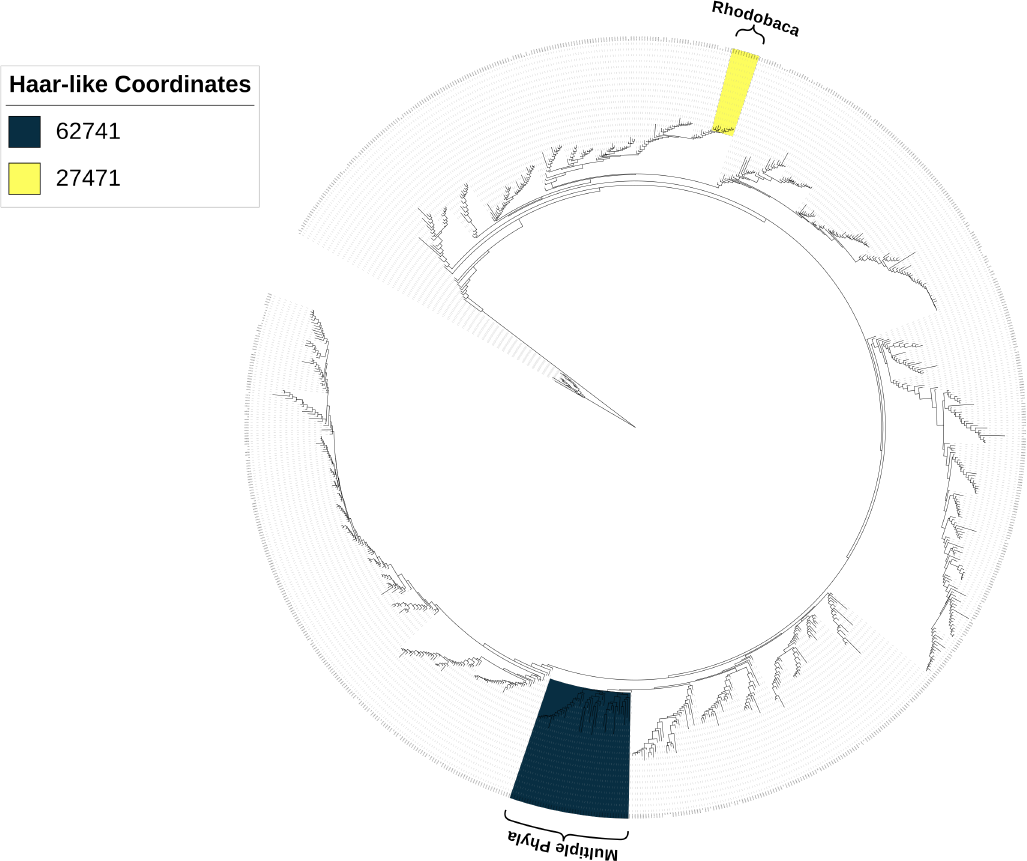
The two most important Haar-like coordinates of the Microbial Mat dataset visualized on Greengenes 97%.

The clade associated with the first dominant Haar-like coordinate (62741) contains OTUs from 15 distinct phyla, making it difficult to link to any taxonomic classification. Notably, the most significant contributors to this coordinate are OTU 156755—an unclassified species within the Nitriliruptoraceae family—and OTU 37747—an unclassified species within the Pseudonocardiaceae family. Referring to Figure 20, we see that both of these OTUs increase in abundance in the deeper mat samples. The relevance of these OTUs in predicting mat depth is consistent with existing literature asserting that Nitriliruptoraceae are haloalkaliphilic and known to thrive in high-pH and high-salinity conditions [63], and Pseudonocardiaceae have been isolated from microbial mats in a lava cave [64].

**Fig 20.**
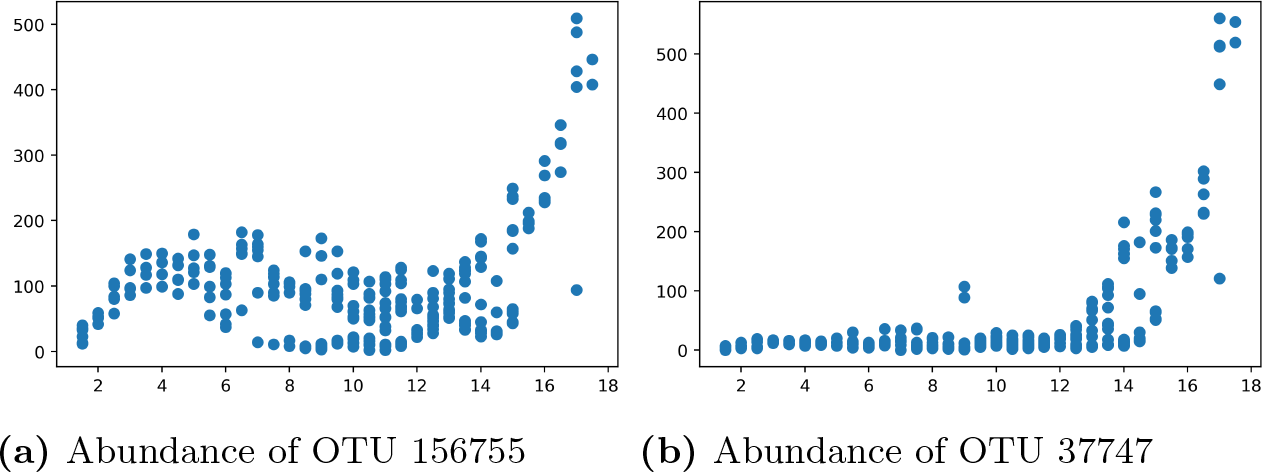
Abundances of the top two contributing OTUs to Haar-like coordinate 62741

**Fig 21.**
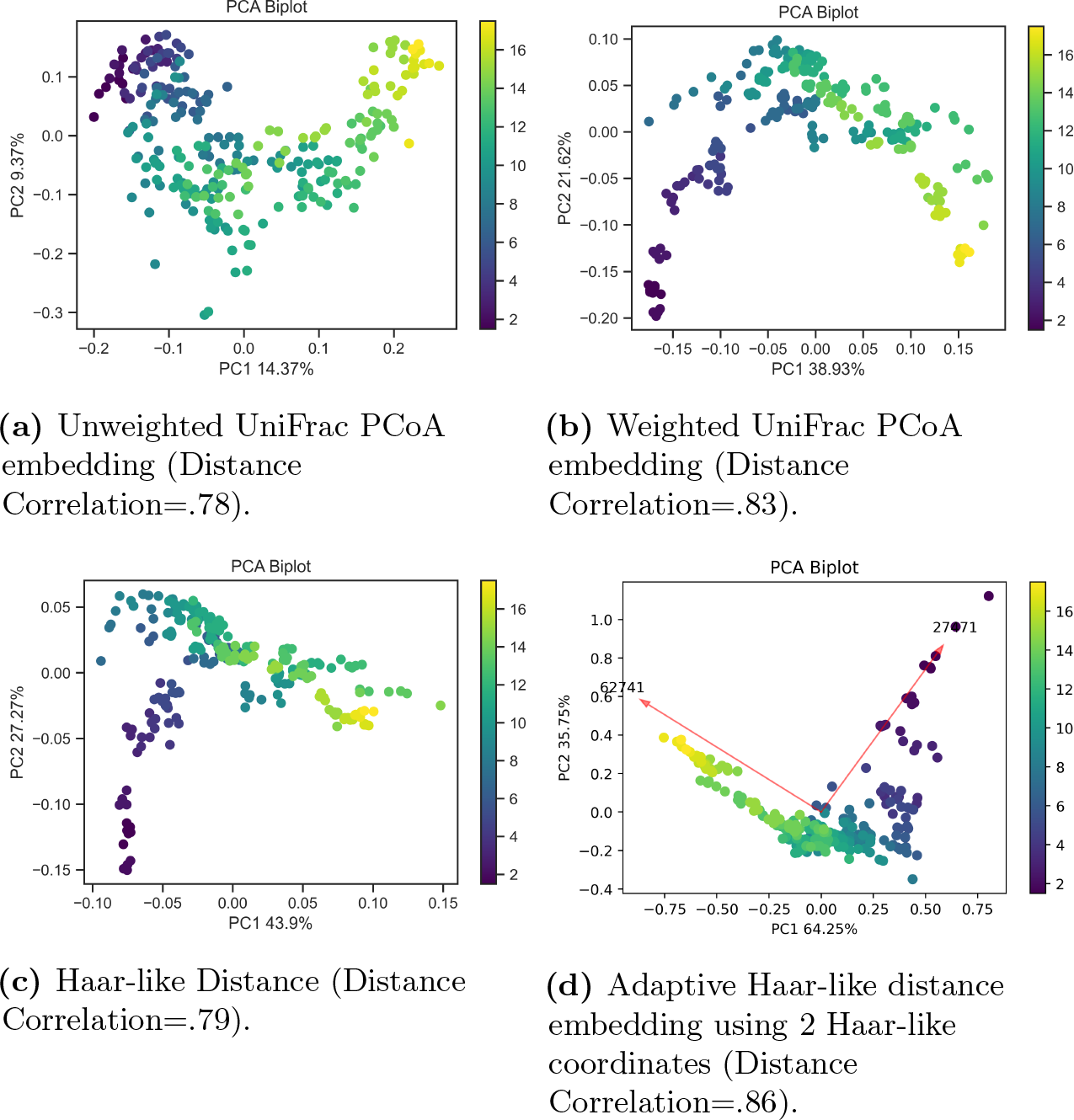
Comparison of the Adaptive Haar-like embedding to unweighted and weighted UniFrac in the Microbial Mat dataset.

In contrast, the second dominant coordinate (27471) consists entirely of the genus Rhodobaca, which are photoheterotrophs that have been previously studied in the context of coastal microbial mats [65].

Despite the lack of connection to established taxonomic classifications, the adaptive Haar-like distance recovers coordinates that are strongly correlated with mat depth (Figure 22). This highlights the utility of our metric as an exploratory tool, particularly in environments with high abundances of uncultured and unclassified species.

**Fig 22.**
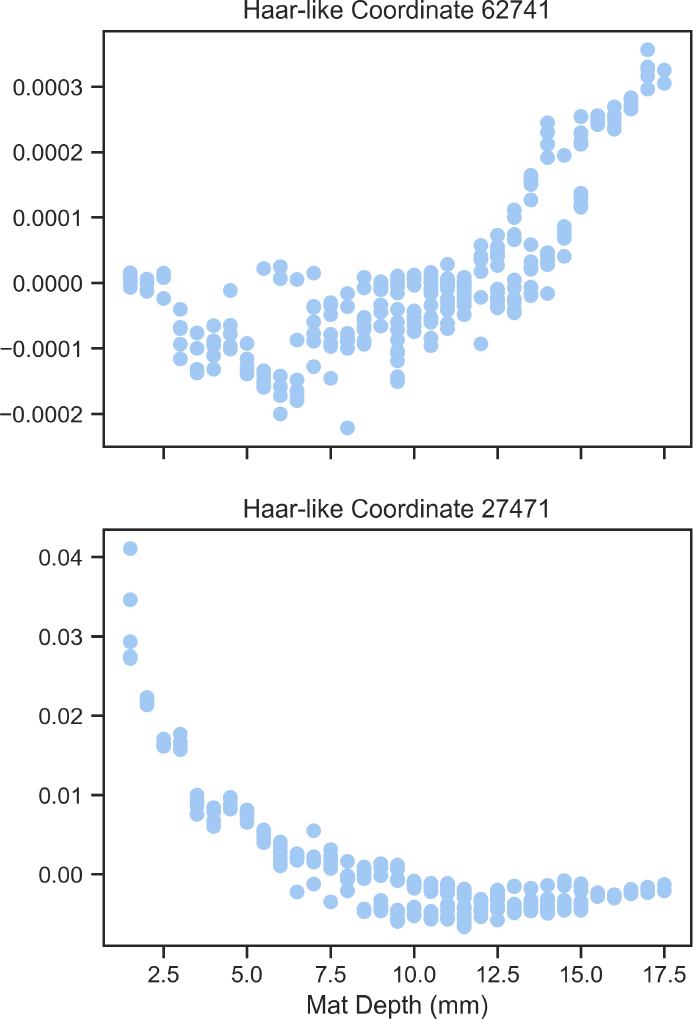
Plots of the top two Haar-like coordinates across samples in the Microbial Mat dataset.

### Model Validation

The preceding section highlighted how the adaptive Haar-like distance can generate insightful embeddings across diverse datasets. However, ensuring that the weights and coordinates derived from our metric can closely match the RF estimates is imperative for the precise categorization of environmental attributes.

Here, we test the Adaptive Haar-like kernel on 16 metagenomic classification problems obtained from the microbiome learning repository (ML Repo) [66], which are listed in Table 1.

**Table 1.** Datasets from ML Repo used in our model comparisons. Acronyms “cd” and “uc” stand for Chron’s disease and ulcerative colitis, respectively.

| Index | Dataset | Task | Sample Type | Number of Samples |
| --- | --- | --- | --- | --- |
| 1 | Montassier 2016 | Bacteremia vs. no bacteremia | Human stool | 28 |
| 2 | David 2014 | Animal vs. plant diet, last diet day | Human stool | 18 |
| 3 | Cho 2012 | Chlortetracycline vs. control, cecal | Mouse cecal contents | 47 |
| 4 | Cho 2012 | Chlortetracycline vs. control, fecal | Mouse pellets | 45 |
| 5 | Cho 2012 | Penicillin vs. vancomycin, cecal | Mouse cecal contents | 47 |
| 6 | Cho 2012 | Penicillin vs. vancomycin, fecal | Mouse pellets | 45 |
| 7 | Gevers 2014 | Control vs. cd, ileum | Ileal biopsies | 140 |
| 8 | Gevers 2014 | Control vs. cd, rectum | Rectal biopsies | 160 |
| 9 | Morgan 2012 | Healthy vs. cd, stool | Human stool | 128 |
| 10 | Morgan 2012 | Healthy vs. uc, stool | Human stool | 128 |
| 11 | HMP 2012 | Male vs. female, stool | Human stool | 180 |
| 12 | HMP 2012 | Stool vs. tongue | Human stool, oral | 404 |
| 13 | HMP 2012 | Subgingival vs. supragingival plaque | Oral | 408 |
| 14 | Yatsunenko 2012 | malawi vs. venezuela | Human stool | 54 |
| 15 | Yatsunenko 2012 | Male vs. female | Human stool | 129 |
| 16 | Kostic 2012 | Healthy vs. tumor biopsy, paired | Colon biopsies | 172 |

To benchmark our method, we compare results to the original RFs and another state-of-the-art interpretable classifier, CoDaCoRe [39], that learns a sparse set of log-ratios to classify metagenomic data. We implemented a stratified 5-fold cross-validation (partitioning the data into 80% training and 20% testing) on each dataset, iterating this process with 5 different randomizations, resulting in 25 unique splits per dataset. In what follows, we outline the precise implementations of each model.

During the training of our adaptive Haar-like kernel, we employ a hyperparameter tuning stage to choose the optimal number of Haar-like coordinates for each dataset. For each randomization of the stratified 5-fold cross-validation, the 80% training data is further split into a training and hyperparameter selection set. The best-performing value of the parameter *s* is then chosen for the final model evaluation on the remaining 20% testing data. The average *s* used for each dataset is reported in Table 2 as “Haar-like Sparsity”.

**Table 2.** Datasets from ML Repo used in our model comparisons. The “taxonomic classification” column lists the lowest taxonomic classification that encompasses all members of the clade corresponding to the given Haar-like coordinate. The phyla in dataset 12 include p Actinobacteria, p Firmicutes, and p Tenericutes

| Index | Dataset | CoDaCore sparsity | Haar-like sparsity | Top Haar-like coordinate | Taxonomic classification |
| --- | --- | --- | --- | --- | --- |
| 1 | Montassier 2016 | n/a | 7.12 | 94036 | o_Clostridiales |
| 2 | David 2014 | n/a | 6.56 | 88556 | o_Clostridiales |
| 3 | Cho 2012 | n/a | 2.16 | 41082 | f_S24-7 |
| 4 | Cho 2012 | n/a | 1.00 | 41082 | f_S24-7 |
| 5 | Cho 2012 | n/a | 2.28 | 41082 | f_S24-7 |
| 6 | Cho 2012 | n/a | 6.56 | 86819 | o_Clostridiales |
| 7 | Gevers 2014 | 2.44 | 7.72 | 86791 | o_Clostridiales |
| 8 | Gevers 2014 | 2.40 | 5.88 | 99273 | f_Lachnospiraceae |
| 9 | Morgan 2012 | 1.76 | 3.16 | 97006 | f_Lachnospiraceae |
| 10 | Morgan 2012 | 1.36 | 4.48 | 98982 | f_Lachnospiraceae |
| 11 | HMP 2012 | 1.96 | 6.80 | 38025 | g_Bacteroides |
| 12 | HMP 2012 | n/a | 5.00 | 99301 | multiple phyla |
| 13 | HMP 2012 | 2.60 | 3.80 | 79173 | g_Parvimonas |
| 14 | Yatsunenko 2012 | n/a | 7.08 | 84112 | f_Ruminococcaceae |
| 15 | Yatsunenko 2012 | 1.88 | 6.84 | 99190 | f_Lachnospiraceae |
| 16 | Kostic 2012 | 1.32 | 7.88 | 98472 | g_Blautia |

We observed better performance in our model by thresholding the Haar affinity matrix in (9). For this, we adopted the popular convention from K-Nearest Neighbors (KNN) classifiers and only kept the weights corresponding to the 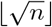 closest neighbors. All other weights were set to zero.

For the RF classifier, we implemented the scikit-learn RF classifier [67] with the default parameter settings.

CoDaCoRe relies on a regularization parameter *λ* to control the trade-off between the sparsity and accuracy of the model. For a fair comparison with our model, we set *λ* = 0 to ensure the highest classification accuracy. The average model sparsity for each dataset is reported in Table 2 as “CoDaCoRe sparsity”. We note that for some datasets, especially those with a small number of samples, CoDaCoRe fails to find a fit due to perfect separation [68]. This occurs during a logistic regression step in CoDaCoRe when an outcome variable entirely segregates a predictor variable, making it impossible to determine a regression coefficient. For these cases, the CoDaCoRe results are omitted (i.e., reported as n/a).

Figures 23 and 24 display boxplots of the accuracy and area under the receiver operating characteristic curve (ROC-AUC score) [69] for the adaptive Haar-like metric, CoDaCoRe, and RF across the sixteen datasets. Table 2 displays additional information about our model, namely the top selected Haar-like coordinate in each dataset and a taxonomic classification that can be associated with the coordinate.

**Fig 23.**
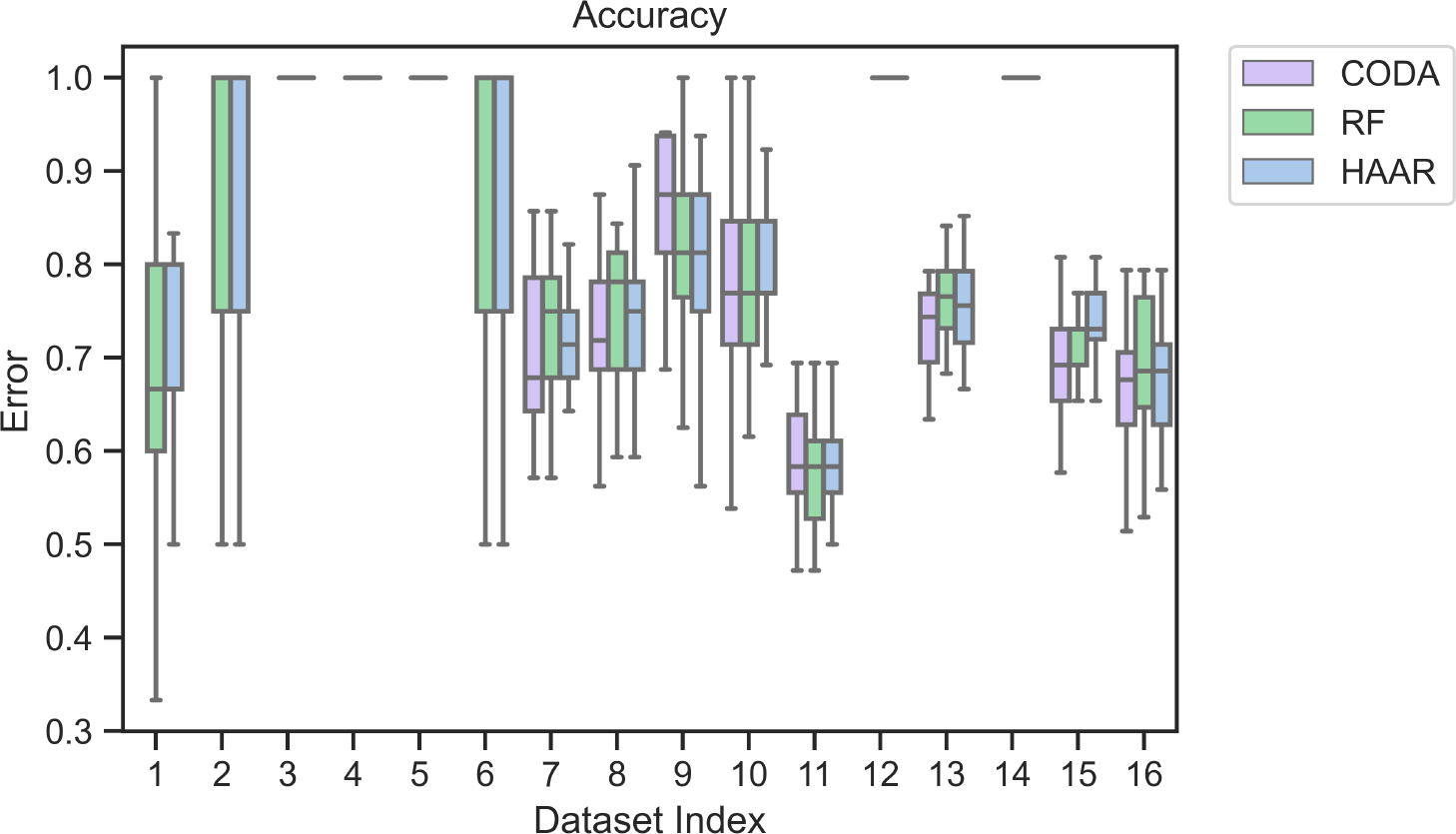
Model accuracy across the 16 ML Repo datasets.

**Fig 24.**
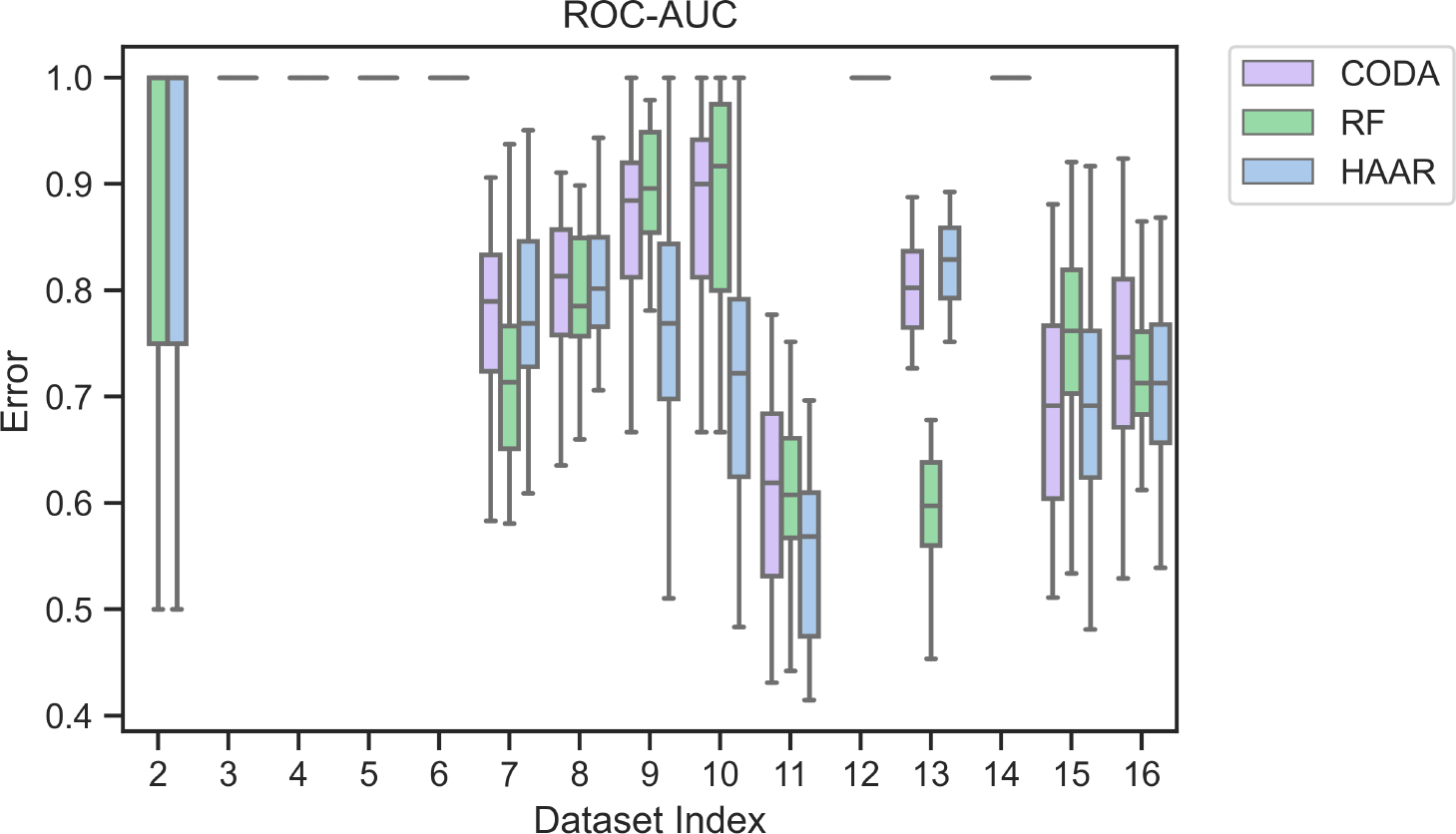
Model ROC-AUC scores across the 16 ML Repo datasets.

Across all the tested datasets, we see that, on average, the three classification models are comparable in terms of accuracy. In terms of AUC, the adaptive Haar-like metric also has comparable performance except on the two datasets from Morgan 2012. Initially, we expected RF to outperform the adaptive Haar-like distance on every dataset because RFs use non-stationary kernels that can fit higher-order interactions than adaptive Haar-like kernels (which are analogous to a Euclidean distance in a modified inner product space). However, because the adaptive Haar-like kernel can only learn linear relationships between microbial abundances, this may reduce over-fitting, allowing for comparable performance to RF in most datasets that we tested.

Interestingly, while CoDaCoRe offers unrestricted OTU selection for crafting log-ratios, the adaptive Haar-like distance restricts OTU groupings based on phylogeny. Although this could enable CoDaCoRe to identify a more concise set of OTUs for model construction, we contend that restricting OTU grouping to the phylogeny enhances biological interpretability.

## Discussion

By learning a metric in a data-dependent manner, the adaptive Haar-like distance can produce accurate and insightful embeddings of metagenomic environments using only a limited number of Haar-like coordinates. The effectiveness of this approach hinges on the interpretation of the Haar-like basis as a wavelet basis on a phylogenetic tree. The sparsity induced by this choice of basis is precisely what allows our algorithm to learn a sparse set of weights that can approximate the far more intricate RF model. Analogous to wavelet denoising [70], selecting only the critical Haar-like coordinates in a dataset and discarding the rest helps build a robust representation of metagenomic environments, thus better differentiating genuine biological signals from noise.

As mentioned earlier, phylofactorization [31] uses a similar coordinate system to decompose microbial abundance in terms of internal nodes of the phylogeny. However, a pivotal distinction in our method lies in its **supervised** approach, where data labels are integrated to discern the most significant clades **within a specific setting**. In contrast, phylofactorization employs an **unsupervised** approach, reminiscent of PCA, to identify clades in the phylogeny that account for the most variance **independently of data labels**.

We underscore that traditional statistical methods employed in Euclidean space do not apply to Haar-like coordinates. This distinguishes our approach from methods like phylofactorization or CoDaCoRe, which utilize isometric log-ratios and have established valid statistical tests [31]. Nonetheless, our method is the only one that exploits phylogenetic structure and takes a supervised learning approach. Further work is therefore necessary to derive statistical tests involving Haar-like coordinates.

Finally, it is worth noting that while we have introduced a data-driven approach for selecting the most significant Haar-like coordinates, our metric can also be applied to investigate user-specified Haar-like coordinate embeddings. In particular, if specific clades hold particular scientific interest, our method can be used to generate biplots, thereby enabling the visualization of the Haar-like coordinates corresponding to those particular clades.

## Conclusion

The adaptive Haar-like distance offers a versatile framework for comparing metagenomic samples from experiments encompassing various biological settings. By tailoring the underlying assumptions to each dataset, our metric learns weights on a reference phylogeny that best differentiate between environmental characteristics of interest. Compared to existing phylogenetic *β*-diversity metrics, the adaptive Haar-like distance can produce quantitatively better embeddings using only a handful of Haar-like coordinates. Our subsequent analysis of the Haar-like coordinates selected in each of the presented datasets confirmed that our metric learning algorithm recovers biologically meaningful splits in the phylogeny. This highlights using our metric as an exploratory tool for uncovering possible relationships between microbial clade abundances and environmental factors. Furthermore, by using the simple adaptive Haar-like kernel to approximate the patterns learned by a more complex but uninterpretable random forest, we offer an interpretable surrogate model with comparable performance.

## Acknowledgments

This work was partially funded by the NSF grant No. 1836914.

